# Chromatin accessibility profiles of castration-resistant prostate cancers reveal novel subtypes and therapeutic vulnerabilities

**DOI:** 10.1101/2020.10.26.355925

**Authors:** Fanying Tang, Shangqian Wang, Chen Khuan Wong, Cindy J. Lee, Sandra Cohen, Jane Park, Corinne E. Hill, Kenneth Eng, Rohan Bareja, Teng Han, Eric Minwei Liu, Ann Palladino, Wei Di, Dong Gao, Wassim Abida, Shaham Beg, Loredana Puca, Michael F. Berger, Anuradha Gopalan, Lukas E. Dow, Juan Miguel Mosquera, Himisha Beltran, Cora N. Sternberg, Ping Chi, Howard I. Scher, Andrea Sboner, Yu Chen, Ekta Khurana

## Abstract

In castration-resistant prostate cancer (CRPC), the loss of androgen receptor (AR)-dependence due to lineage plasticity, which has become more prevalent, leads to clinically highly aggressive tumors with few therapeutic options and is mechanistically poorly defined. To identify the master transcription factors (TFs) of CRPC in a subtype-specific manner, we derived and collected 29 metastatic human prostate cancer organoids and cell lines, and generated ATAC-seq, RNA-seq and DNA sequencing data. We identified four subtypes and their master TFs using novel computational algorithms: AR-dependent; Wnt-dependent, driven by TCF; neuroendocrine, driven by ASCL1 and NEUROD1 and stem cell-like (SCL), driven by the AP-1 family. The transcriptomic signatures of these four subtypes enabled the classification of 370 patients. We find that AP-1 co-operates with the inhibitable YAP/TAZ/TEAD pathway in the SCL subtype, the second most common group of CRPC tumors after AR-dependent. Together, this molecular classification reveals new drug targets and can potentially guide therapeutic decisions.

## Main Text

Untreated prostate cancers rely on androgen receptor (AR) signaling for growth and survival, forming the basis for the nearly universal initial efficacy of androgen deprivation therapy (ADT). Yet, the disease usually relapses and progresses to a lethal stage termed castration-resistant prostate cancer (CRPC). Reactivation of AR signaling represents the most common mechanism of CRPC and this understanding has led to the development of highly potent next generation AR inhibitors which are now frequently administered upfront with ADT(*1*). However, the contemporary highly potent repression of AR signaling also exerts selective pressure to generate AR-independent tumors that rely on distinct survival mechanisms. This transition away from AR-dependence frequently accompanies a change in phenotype that resembles developmental trans-differentiation or ‘lineage plasticity’ (*2*). While neuroendocrine prostate cancer (NEPC) is the most studied type of lineage plasticity (*3*), recent studies have shown that most AR-null tumors do not exhibit neuroendocrine features. These tumors have been termed “double-negative prostate cancer” (DNPC) and their drivers are poorly defined (*4, 5*).

Mechanistic studies in CRPC are limited by the lack of genetically defined patient-derived models that recapitulate the disease spectrum. To overcome this, we have developed a biobank of organoids from patient biopsies that provide a unique resource to study the landscape of metastatic CRPC and allow for testing predictions by functional validation assays. These organoids recapitulate the phenotypic diversity of CRPC, including AR-positive adenocarcinoma, neuroendocrine/small cell carcinoma and AR-negative (neg) adenocarcinoma. They also represent a wide spectrum of genotypic alterations, including AR amplification, ERG fusion, RB1 deletion and alterations of components in multiple signaling pathways (*6, 7*).

Transcription factors (TFs) are drivers of tumor lineage plasticity (*8*). To identify CRPC subtypes and predict their master TFs, we developed a novel ranking algorithm that combines three layers of information at the transcriptomic, epigenetic and regulatory network levels. We generated high-quality ATAC-seq and RNA-seq data for 24 metastatic prostate cancer organoids and cell lines, and included five additional samples with published matching datasets (*9*) in the downstream analysis. The 29 samples together represent the largest collection of CRPC in vitro models, and the dataset is also the largest catalogue of ATAC-seq profiles for metastatic prostate cancer to our knowledge. We classified the 29 CRPC models into four subtypes and predicted corresponding master TFs of each subtype. The analysis revealed two subtypes of AR-neg or low (neg/low) tumors that have not been reported before, one driven by the AP-1 family of TFs and another by TCF/LEF of the Wnt signaling pathway. Additional data analysis and experimental results indicate that AP-1 works together with YAP, TAZ and TEAD, revealing YAP/TAZ as actionable targets in one subtype. The RNA-seq signatures derived from the organoids enable the classification of 370 patient samples from two independent CRPC datasets into the four subtypes, suggesting that our results can be directly used for patient stratification for appropriate therapeutic opportunities.

## Results

### Biobank of patient-derived organoids of metastatic CRPC

For this work, we generated and characterized 10 novel prostate cancer organoids from specimens of patients with metastatic CRPC (MSKPCa8-MSKPCa17) to add to our biobank (*6, 7*). In general, the organoids were from patients with aggressive disease, with short response to initial ADT and rapid progression on next-line AR inhibition therapy (table S1). In culture, the organoids adopted similar histology as the tissue (fig. S1A and S1B) and the neuroendocrine samples maintained IHC staining of synaptophysin (SYP) (fig S1C).

We generated mutational and copy number profiles of each organoid as well as 7 of 10 matching tumor biopsy specimens using MSK-IMPACT. The copy number landscape was similar between tumors and organoids, and was representative of metastatic prostate cancer when compared to the Stand Up to Cancer (SU2C) cohort (*10*) (Fig. 1A). We observed a mean of 3.42 somatic mutations per patient, which is similar to the entire cohort of metastatic prostate cancer patients sequenced by MSK-IMPACT (*11*) (Fig. 1B). The majority of organoids exhibit exactly the same copy number variations (CNVs) and single nucleotide variants (SNVs) as the original biopsies (fig. S1D, table S2 and S3). In fact, organoids represent higher purity than the original biopsies as shown by the increased tumor allelic frequency of SNVs and CNVs (Fig. 1A and 1B).

**Fig. 1.**
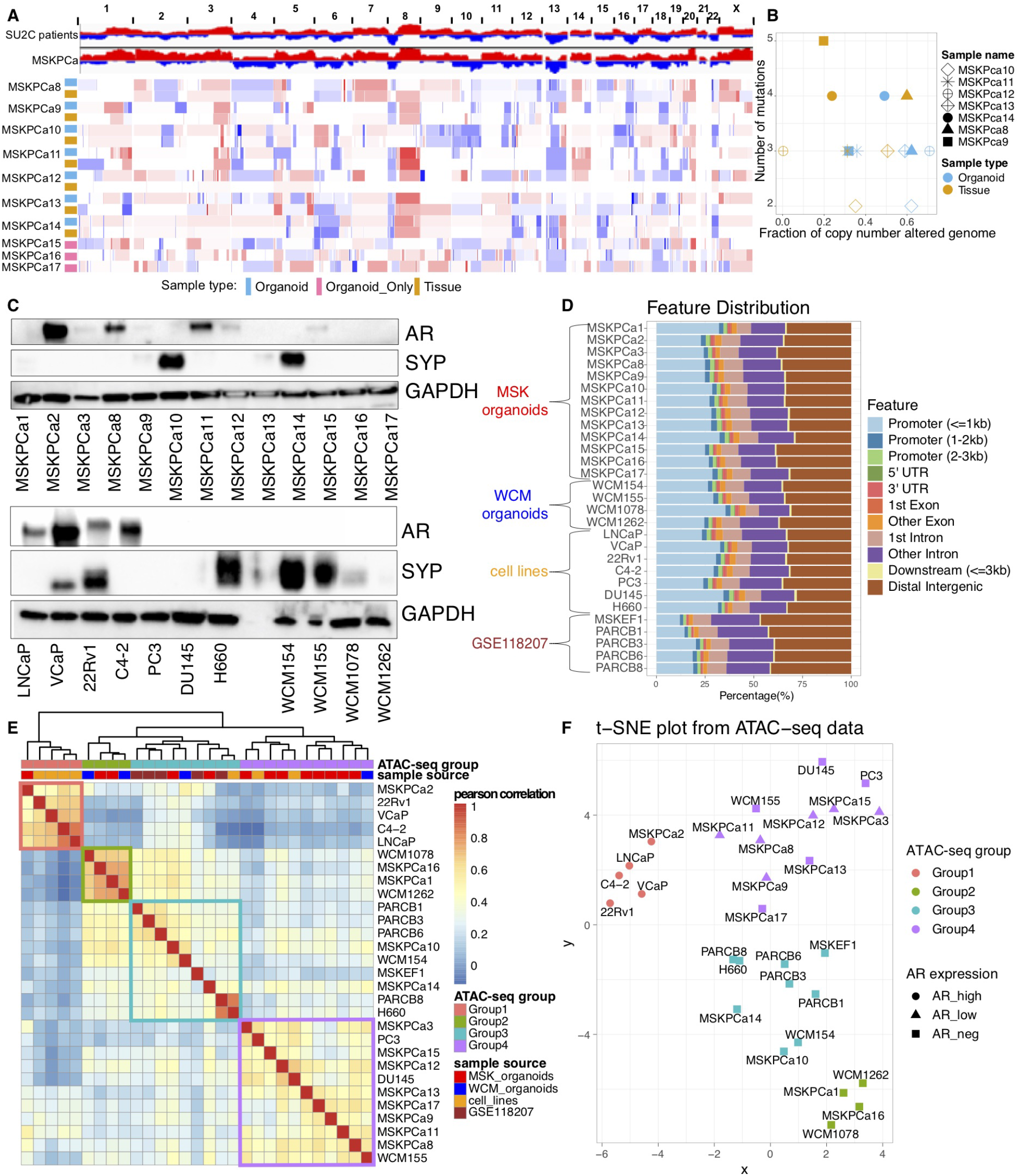
Characterization of novel patient-derived organoids from metastatic CRPC and the classification of metastatic prostate cancer into four molecular subtypes based on chromatin accessibility. (**A**) Top: Significant genomic aberrations in the prostate onco-genome from the SU2C CRPC patient samples (*10*) and MSKPCa organoids. Bottom: Copy number landscape of the ten patient derived organoid lines and seven matching tumor tissues using IMPACT sequencing data. Shades of red and blue represent levels of gain and loss. (**B**) Number of mutations and fraction of copy number altered genome of the seven organoids and their matching patient tumor tissues. (**C**) Immunoblot showing the expression of AR and SYP across the 24 organoids and cell lines. (**D**) Feature distribution of the mapped ATAC-seq peaks across all samples. (**E**) Correlation heatmap based on the normalized number of reads of the top 1% variable peaks across all samples. The ATAC-seq group and the sample source are indicated for each sample. The colors of the four ATAC-seq groups are kept consistent throughout the paper. (**F**) Unsupervised t-SNE on the top 1% variable accessible peaks across all samples.

### Chromatin accessibility landscape reveals four molecular subtypes of metastatic prostate cancer

We performed ATAC-seq assays for 24 metastatic prostate cancer models, including 17 patient-derived organoids and 7 cell lines with two biological replicates for each (fig. S2A to S2E, table S4). These patients were chosen to represent the diversity of CRPC and include 16 samples that are AR-null or AR-low tumors. We also included ATAC-seq data from 5 NEPC models generated by Lee JK *et al* (*9*) (Fig. 1D). Overall, we identified 848,913 reproducible peaks (irreproducible Discovery Rate < 0.05 between biological replicates and observed in more than one sample). The majority of the ATAC-seq peaks map to distal intergenic and intronic regions, similar to data reported by other groups (*12*) (Fig. 1D). We identified four CRPC subtypes using consensus k-means clustering on the regions showing most variable accessibility (fig. S2F-S2H). We obtained the same four groups using other clustering approaches, such as hierarchical clustering and t-SNE (Fig. 1E and 1F). There is no significant difference between the number of peaks among the four subtypes (fig S2I). One of the four subtypes consists of high AR expressing cells, including LNCaP, VCaP, 22Rv1, C4-2 and MSKPCa2 (Fig. 1C). The promoter region of *KLK2,* which is an AR target gene, is also more accessible in this subtype relative to others (fig. S2J). Another subtype includes published NEPC samples, i.e. H660, WCM154, PARCB1, PARCB3, PARCB6, PARCB8, MSKEF1 (a derivative of published neuroendocrine organoid MSKPCa4), and two unpublished organoid lines, MSKPCa10 and MSKPCa14, with high SYP expression (Fig. 1C) and small-cell carcinoma phenotype (fig. S1A and S1C). The remaining two subtypes are enriched with NE-negative, AR-neg/low samples (Fig. 1C). For further biological characterization, we integrated the ATAC-seq data with RNA-seq and DNA sequencing data.

### Transcriptomic profiles of the four CRPC subtypes

We next analyzed the transcriptomes of the 24 samples using t-SNE and found that the clusters agree with the subtypes identified using ATAC-seq (Fig. 2A). Based on gene set enrichment analysis (GSEA) and selective marker gene expression, we named the four subtypes as follows: 1) Group1_AR, which is enriched in AR signature (*13*) (Fig. 2B, 2F and 2G, fig. S3B). 2) Group2_WNT, which is enriched in Wnt signaling, and includes WCM1078, WCM1262, MSKPCa1 and MSKPCa16 (Fig. 2C, 2F and 2G, fig. S3B). 3) Group3_NEPC samples are strongly enriched in the NEPC signature (*14*), in agreement with the pathology classification of NE (fig. S1A and S1C) and have high expression of NEPC markers, including SYP, CHGA and DLL3 (Fig. 2D, 2F and 2G, fig. S3B). 4) Group4_SCL consists of stem cell-like (SCL) tumor cells, including 9 organoids and prostate cancer cell lines DU145 and PC3.

Group4_SCL has not been previously defined. Samples in this subtype are enriched with the mammary stem cell signature, with high expression of cancer stem cell markers CD44 and TACSTD2 (TROP2A) (Fig. 2E to 2G, fig. S3B). Samples in Group4 are also enriched with multiple hallmark pathways, including TGF-beta, epithelial-mesenchymal transition, inflammation, TNF-alpha signaling and interferon responses (fig. S3A).

**Fig. 2.**
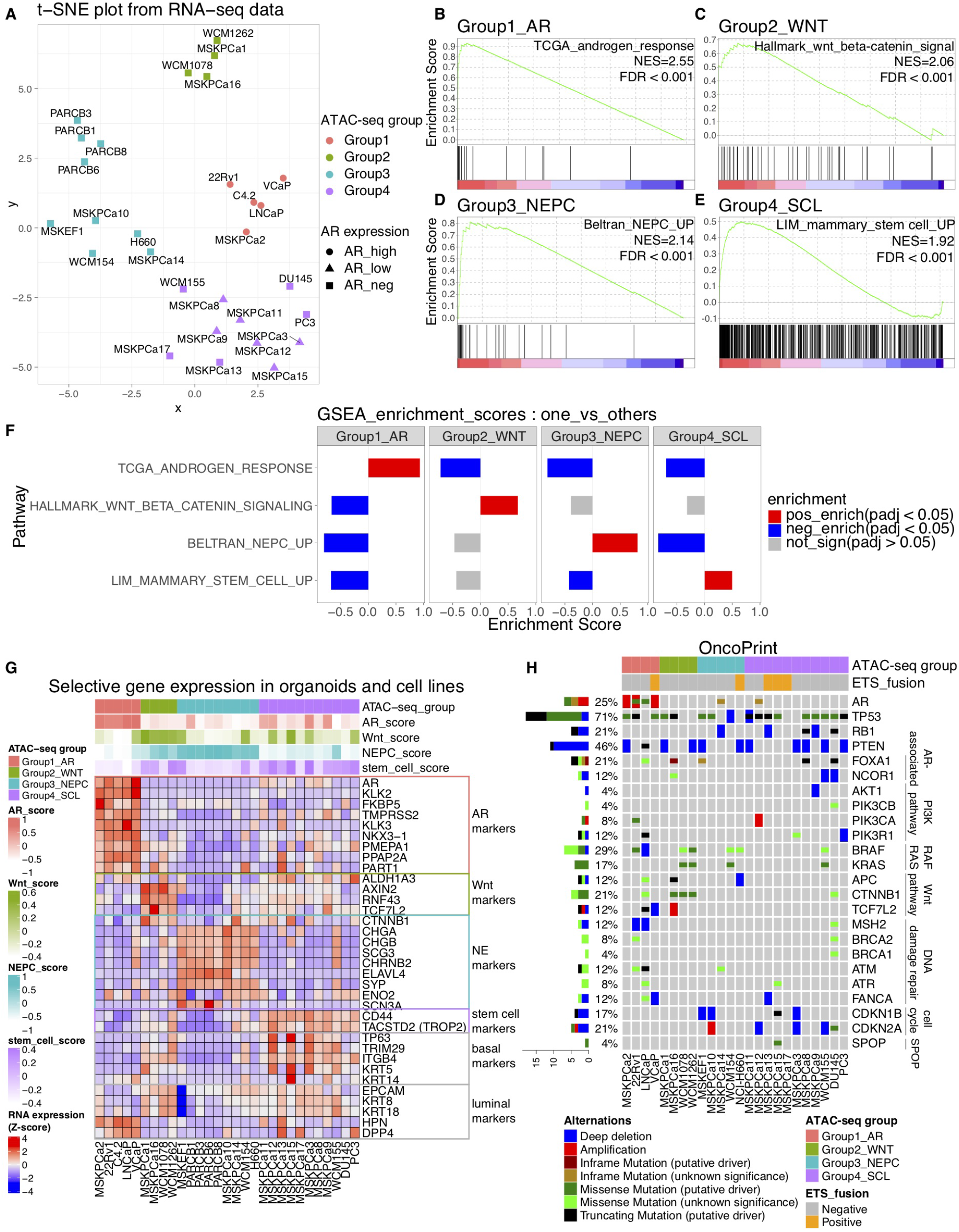
Transcriptomic and genomic characterization of the four CRPC subtypes defined by ATAC-seq. (**A**) Unsupervised t-SNE on the mRNA expression values of the 1,000 most variably expressed protein-coding genes across all samples. (**B-F**) GSEAs show representative enriched pathways in Group1_AR (**B**), Group2_WNT (**C**), Group3_NEPC (**D**) and Group4_SCL (**E**). Enrichment score and p-values indicate the four signals are significantly positively enriched in corresponding subtypes but not in others (**F**). (**G**) Heatmap shows the relative expression of subtype-specific marker genes and basal/luminal genes across all organoids and cell lines. The ATAC-seq group and signature scores of the four representative pathways for each sample are shown at the top. (**H**) Oncoprint shows the genomic alterations of the 24 samples with DNA-sequencing data. MSKPCa8~17 were sequenced with MSK-IMPACT. The SNVs and CNVs for cell lines (LNCaP, 22Rv1, VCaP, H660, DU145 and PC3) were collected from CCLE. Whole-exome sequencing (WES) was performed for other samples.

Quantitative PCR (qPCR) of the selective marker genes (NKX3-1, TMPRSS2, AXIN2, RNF43 and SYP) across all samples confirm the four subtypes defined above (fig. S11).

We find that compared with Group1_AR, the other three groups are enriched with basal signature (*15*) and prostate basal stem cell signature (*16*), with Group4_SCL having the highest enrichment score (fig. S3C) and expression of basal cell markers (Fig. 2G). Group2_WNT and Group3_NEPC, but not Group4_SCL, have significant depletion of luminal signature compared to Group1_AR (fig. S3C). In addition, consistent with previous studies of AR-neg/low tumors, Group2_WNT and Group4_SCL show strong enrichment of FGFR signaling and expression of selective FGF ligands and receptors compared to the other two groups (fig. S4A and S4B) (*4*).

### Genomic characterization and loss of tumor suppressors in the four CRPC subtypes

All samples in Group1_AR have AR amplification and/or AR mutation (Fig. 2H), in agreement with the high AR activity in the group. In Group2_WNT, all the four samples show alterations in the Wnt signaling pathway (*17*) (Fig. 2H and fig. S4C). Three samples show hot spot mutations in CTNNB1 (beta-catenin), the intracellular signal transducer of Wnt signaling, and the event is observed exclusively in Group2_WNT (fig. 2H and fig. S4C). The fourth sample has shallow deletion of the Wnt signaling negative regulator APC and gain of the activator RSPO2 (*17*) (fig. S4C).

Loss of tumor suppressors TP53, PTEN, and RB1 is associated with lineage plasticity and aggressive disease in CRPC (*15, 18*). We find that TP53 is the most frequently mutated gene, with putative driver mutations or deep deletions in 17 samples across all four groups. RB1 and PTEN have bi-allelic alterations in 4 and 10 samples respectively (Fig. 2H and fig. S4D). Using RNA-seq and immunoblot analysis, we find that an additional 7 samples exhibit RB1 loss and 4 samples exhibit PTEN loss (fig. S4E and S5, table S5). Overall, we find an enrichment of RB1 loss in AR neg/low samples (11/20) compared to Group1_AR (0/5) (p-value=0.03768, one-tail Fisher’s exact test) while there is no statistical difference in PTEN and TP53 alterations between Group1_AR and others. AR-independent CRPC have worse prognosis and thus these results agree with the recent whole-genome sequencing study indicating that samples harboring bi-allelic RB1 alterations are associated with shorter survival, while there is no prognostic difference for patients with alterations at PTEN and TP53 (*19*). It is notable that while 9/20 lines exhibit loss of both TP53 and RB1 in the AR-neg/low samples, only two are overtly NEPC (fig. S1A, S1B and S1C), consistent with recent observations that loss of TP53 and RB1 in prostate carcinoma attenuates AR signaling but does not uniformly induce neuroendocrine phenotype (*20*). This further highlights the importance of transcriptomic and epigenetic analysis in defining CRPC subtypes.

### Construction of regulatory networks and identification of master TFs

To identify master TFs that drive the subtype-specific transcriptome, we first identified the hubs in regulatory networks that target more genes in a given sample. We constructed regulatory networks by integrating of ATAC-seq and RNA-seq data, and built the peak-gene links based on the correlation between chromatin accessibility at ATAC-seq peaks and expression of genes within +/− 0.5Mb (Fig. 3A, step1) (*12*). In total, we identified at least one peak-gene link for 4752 protein-coding genes (table S6). 75.2% of the peaks are predicted to regulate only one gene, and on average the expression of one gene is correlated with the activity of three peaks (Fig. 3B and 3C). To uncover TF-DNA binding sites in the accessible regions, we used a footprinting method called HINT-ATAC and a curated collection of sequence binding motifs for 809 TFs from CisBP (Fig. 3A, step2). HINT-ATAC searches for regions with depletion of transposition due to protection from TF binding in otherwise nucleosome-free DNA (*21*). By combining peak-gene and TF-peak links, we constructed TF-gene links and generated the regulatory network for each sample (Fig. 3A, step3 and 4). We define TF out-degree as the number of target genes it regulates in the network (fig. S6).

**Fig. 3.**
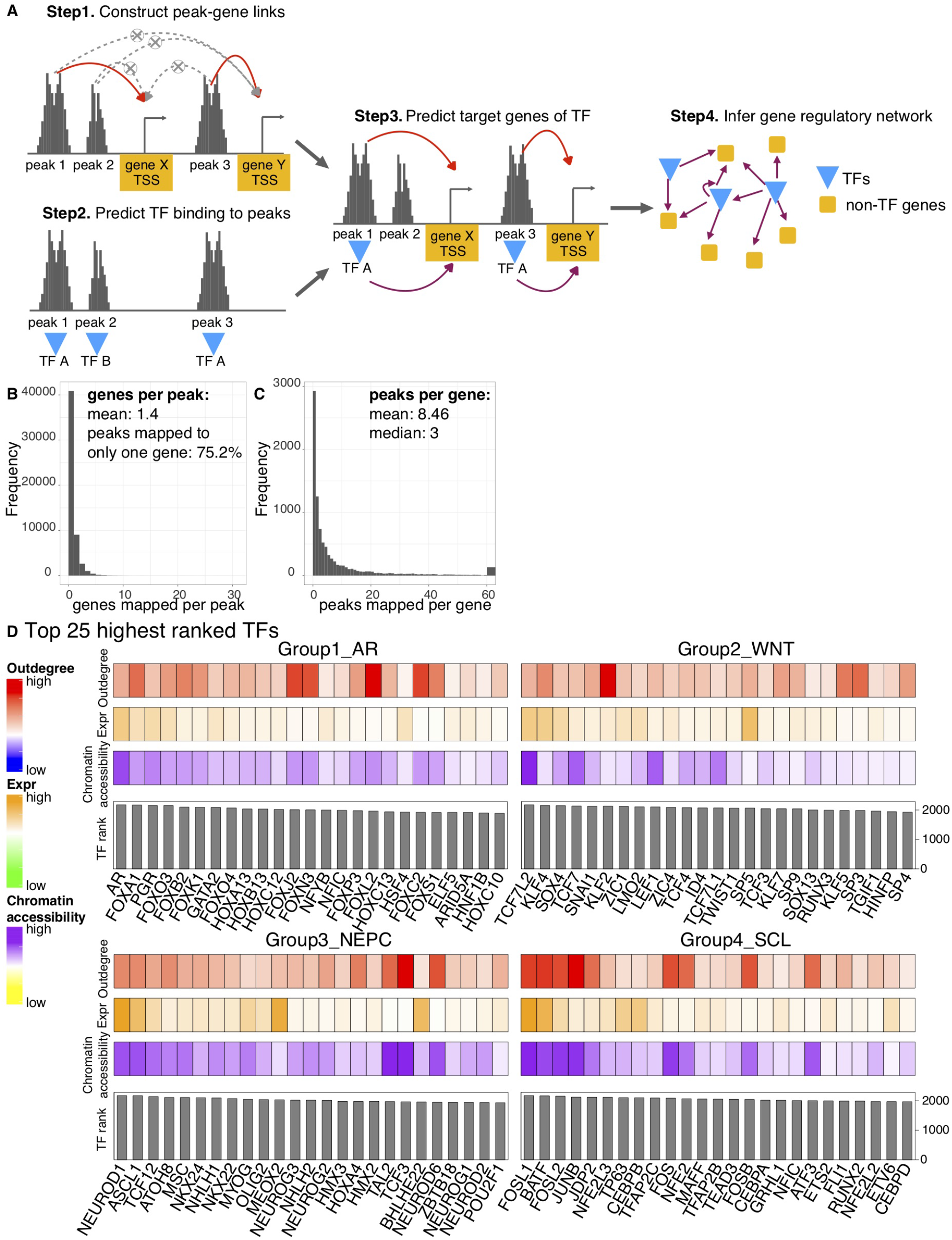
Identification of the master transcription factors (TFs) of each subtype. (**A**) Schematic illustrating the construction of sample-specific regulatory networks using ATAC-seq and RNA-seq data. (**B**) Distribution of the number of genes linked per peak. **(C**) Distribution of the number of peaks linked per gene. (**D**) Rank order of the potential master TFs for each of the four subtypes (top 25 are shown). For each TF, the relative contributions of 3 metrics to TF rank are shown (Expr, expression). In Group4_SCL, FOSL1, BATF, FOSL1, JUNB, JDP2, FOS, MAFF, FOSB and ATF3 belong to the AP-1 family.

Next, we identified master TFs for each subtype as those at the top of the gene regulation hierarchy (Fig. 3D, table S7). Essentially these are the TF hubs in regulatory networks with high expression and motif enrichment at open chromatin regions. Thus, each TF is ranked based on a combination of three metrics: 1) its differential outdegree: *O_diff* (fig. S7A); 2) its differential chromatin accessibility at its motifs: *A_diff* (fig. S7B); and 3) its differential gene expression: *E_diff* in a given subtype relative to others (Fig. 3D and fig. S8).

AR plays a pivotal role in the survival and proliferation of AR-dependent prostate cancer and it is indeed identified as the top candidate in Group1_AR using our approach. The second candidate identified by our approach in this subtype is FOXA1, a forkhead family TF, which has been shown to function as a pioneering factor, increase chromatin accessibility and recruit AR to prostate-specific enhancers (*22, 23*). In Group3_NEPC, the top two TFs are neurogenic differentiation factor 1 (NEUROD1) and achaete-scute homologue 1 (ASCL1). NEPC and small cell lung cancer (SCLC) have been shown to be similar at phenotypic and molecular level, and ASCL1 and NEUROD1 have been demonstrated to be the main drivers in SCLC (*9, 24*).

In Group2_WNT Transcription Factor 7 Like 2 (TCF7L2) gets the highest rank. Also known as TCF-4, TCF7L2 has been shown to be the key driver in colorectal cancer upon upstream Wnt pathway gene alterations such as APC mutations (*25*). Other TCF and LEF TFs are also listed as top candidates, including LEF1/LEF, TCF7/TCF-1 and TCF7L1/TCF-3. Upon Wnt pathway activation, TCF and LEF work together with beta-catenin in the nucleus and function as activators to promote the expression of downstream genes (*25*). Thus, the predicted master TFs in Group2_WNT match well with the strong Wnt signaling activation.

In Group4_SCL, we identify the AP-1 family as the top master TF candidates, with FOSL1 having the highest rank. AP-1 is a TF complex assembled through homo- or hetero-dimerization of members of the Fos and Jun family (*26*). The Fos family includes FOSL1, FOSL2, FOS, FOSB, while the Jun family includes JUN, JUNB and JUND. AP-1 has been shown to be activated by multiple upstream signals including growth factors, hormones, cytokines, inflammation and stresses. It controls the expression of many downstream genes related to cell division, apoptosis, cell migration and immunity (*26*).

Besides subtype-specific manner, we also predicted the important TFs in a sample-specific manner. To do this, we analyzed all TFs’ relative expression and chromatin accessibility at their motifs for all samples in Groups 2 to 4 and compared it to the average of Group1_AR samples. This analysis shows that the master regulators identified in a subtype-specific manner agree with the sample-specific results (fig. S9). We also find significant correlation between the expression of master TFs and accessibility at their motifs suggesting they likely exhibit pioneering activity (fig. S10) (*12*). Finally, the relative expression levels of predicted master TFs in each of the four groups were confirmed by qPCR tests across all samples (fig. S11).

### Classification of CRPC patients using transcriptomic signatures for the four subtypes

Next, we used RNA-seq data from 370 CRPC patients to assign each patient to the four subtypes (*27*). We derived the signature genes for each of the four subtypes as the ones with higher expression in one group relative to others in organoids and cell lines and filtered out genes with low expression or low variance in CRPC patient samples (table S8). A template was constructed for each subtype by combining all the signature genes and assigning a value of 1 to the signature genes of that group, while 0 was assigned to the signature genes of other groups. To assign the CRPC patient samples to a subtype, we used Nearest Template Prediction (NTP) algorithm (*28, 29*), which involves computing the cosine distance (*d*) between each patient’s RNA-seq data and each of the four templates and estimating the statistical significance by random re-sampling (Fig. 4A).

**Fig. 4.**
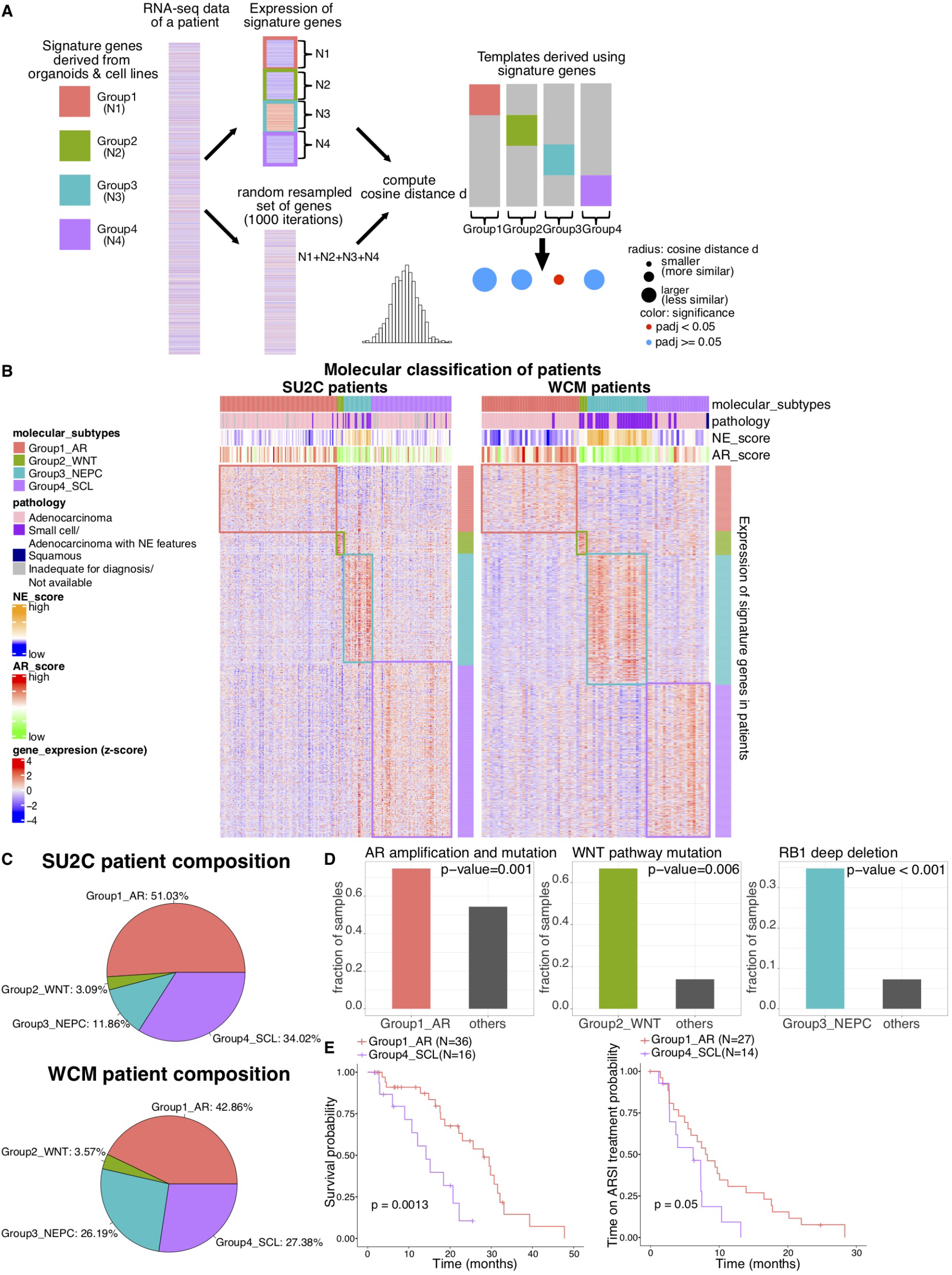
Classification of CRPC patients using transcriptomic signatures for the four subtypes. (**A**) Schematic illustrating the assignment of each patient to one of the 4 groups using nearest template prediction (See Methods for details). (**B**) Heatmaps showing relative expression of signature genes in SU2C and WCM patients. Top annotations indicate the AR score, NE score, pathology classification and molecular subtypes of each patient. (**C**) SU2C and WCM cohorts show similar patient compositions. (**D**) AR amplification and mutation, WNT pathway component mutations and RB1 deep deletion are enriched in Group1_AR, Group2_WNT and Group3_NEPC SU2C patients respectively. (**E**) Group4_SCL patients have significantly worse overall survival and a tendency of shorter time on ARSI treatment compared to Group1_AR SU2C patients.

We applied the method to two cohorts of CRPC patients with polyA-enriched RNA-seq data, including 270 published SU2C patients (*27*) and 100 in-house patients sequenced at Weill Cornell Medicine (WCM) (Fig. 4B). The majority of the patients (194/270 SU2C and 84/100 WCM) were assigned to one of the four subtypes (fig. S12 and S13, table S9). The relative ratios of the four subtypes of patients are similar between cohorts, with the largest group being Group1_AR (having around half of the patients), then Group4_SCL, followed by Group3_NEPC and Group2_WNT. For WCM1078, WCM1262, WCM155 and MSKPCa2 organoids, RNA-seq data are available from matching tumor samples, which are assigned to the same subtype as the organoids (table S9).

The genomic alterations, marker gene expression and pathology analysis provide validation of patient classification. Group1_AR patients show enrichment of AR amplification (Fig. 4D and fig. S15A) and have higher AR expression and AR score (fig. S14) compared to other groups. Group3_NEPC patients have higher SYP expression and NE score (*27*) (fig. S14) compared to others, and are enriched with RB1 deep deletion (fig. 4D and fig. S15A). The majority of patients in this class were also diagnosed as having either small cell, NEPC, or adenocarcinoma with NE features based on histology analysis (Fig. 4B). Patients in Group2_WNT show elevated expression of AXIN2 (fig. S14B), and an enrichment of mutations of Wnt pathway components (Fig. 4D, fig. S15A and S15B). We observe increased expression of the stem cell marker CD44 in Group4_SCL patients (fig. S14B) compared to others as expected, but no consistent enrichment of gene or pathway alterations at the genomic level (fig. S15A, fig. S15C).

Among the 270 SU2C patients, 80 of them have corresponding overall survival time, and 61 of them have drug treatment time when treated with next generation androgen signaling inhibitors (ARSIs) enzalutamide and abiraterone acetate. We find Group4_SCL is associated with worse overall survival compared to Group1_AR patients, and shows a tendency of shorter time on ARSI treatment using cox log-rank statistics (Fig. 4E), indicating that the ARSI treatments were less effective for Group4_SCL patients. We could not compare the patient survival of Group1_AR to other subtypes because there were fewer than 5 samples for Groups 2 and 3 (table S10).

### AP-1 cooperates with YAP, TAZ and TEAD in Group4_SCL

The proportion of patients classified as Group4_SCL is the second largest among both SU2C (34.02%) and WCM (27.38%) cohorts (Fig. 4C), thus we further explored samples in this subtype. We focused on DU145, an AR-negative cell line and MSKPCa3, an AR-low organoid as Group4_SCL models for experimental validations.

#### Cell competition assays for AP-1

We identified AP-1 family, in particular FOSL1, as the top candidate master TF for Group4_SCL (Fig. 3D). Experimental measurement of the expression of various AP-1 components across the four subtypes confirms FOSL1 as the AP-1 gene with highest relative expression in Group4_SCL samples compared to others (fig. S16A) while it is barely detectable in Group1_AR samples at mRNA and protein levels (fig. S16A and S16B). To directly assess the importance of FOSL1 on tumor growth, we performed cell competition assays in DU145 and MSKPCa3, with an AR-dependent organoid MSKPCa2 as the negative control. We transduced the cells with constructs containing GFP, Cas9 and sgRNAs against FOSL1, RPA3 (positive control) or Rosa26 (negative control) and monitored the relative proportion of GFP-positive sgRNA-expressing cells compared with GFP-negative cells over time by fluorescence-activated cell sorting (fig. S16C). A strong depletion of GFP positive sgFOSL1 was observed in both DU145 and MSKPCa3, but not in MSKPCa2, supporting our prediction that FOSL1 is important for tumor progression in Group4_SCL (Fig. 5A, fig. S16D and 16E).

**Fig. 5.**
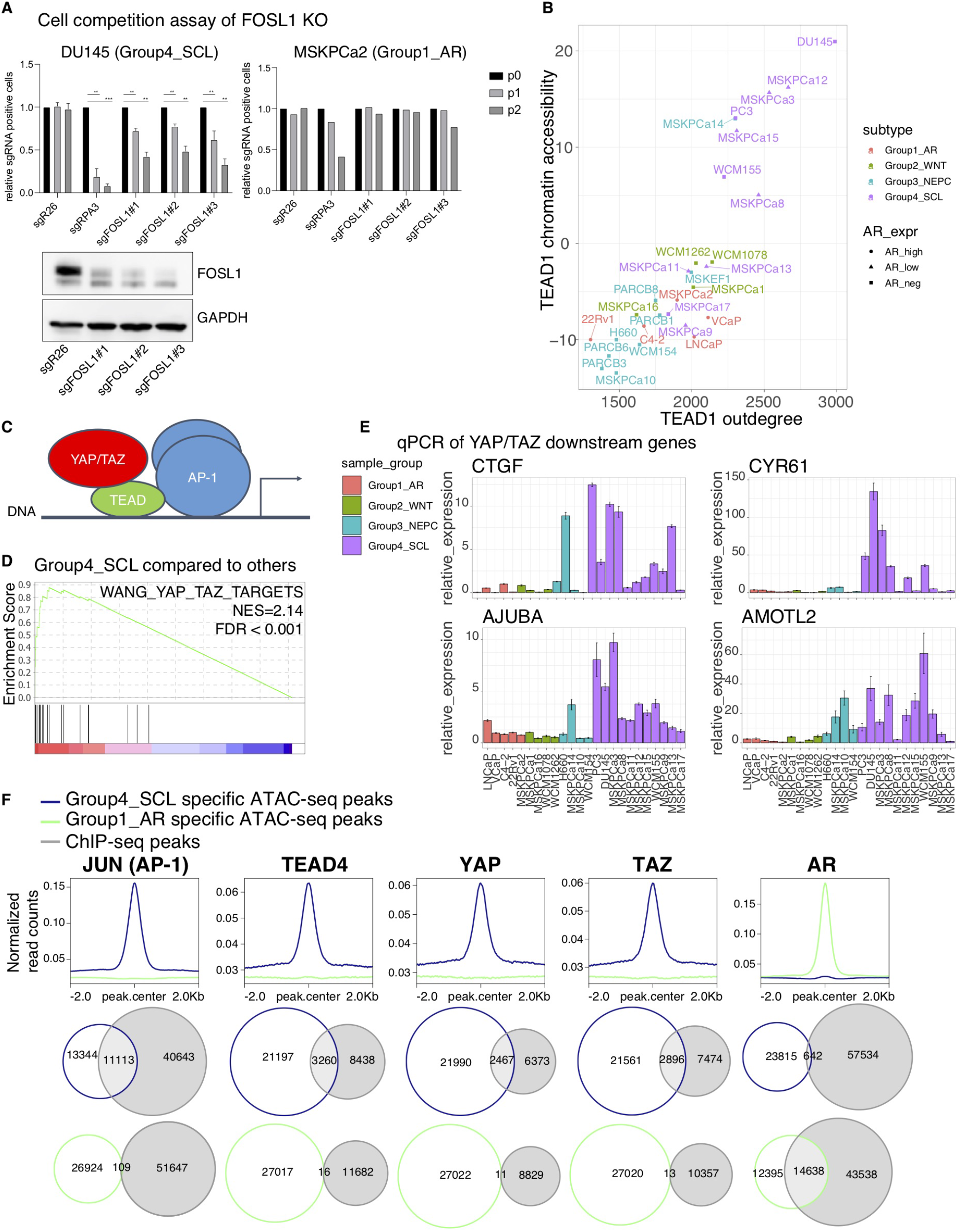
AP-1 works together with YAP, TAZ and TEAD in Group4_SCL. (**A**) Percentage of GFP positive DU145 (left) or MSKPCa2 (right) expressing CRISPR guides against FOSL1 or sgR26 (negative control) or sgRPA3 (positive control) over 3 passages. Mean ± SD. Two-tailed unpaired t-test, n=2 for DU145, n=1 for MSKPCa2. Knockout of FOSL1 was confirmed in DU145 by Western blot. (**B**) Group4_SCL is enriched with samples of high TEAD1 out-degree (from the constructed regulatory networks) and high chromatin accessibility (z-score from ChromVAR) at TEAD1 motifs. (**C**) Schematic showing the co-operation of AP-1 with YAP/TAZ and TEAD in Group4_SCL samples. (**D**) GSEA plot showing that YAP/TAZ signature is strongly enriched in Group4_SCL organoids and cell lines compared to other samples. (**E**) qPCR of YAP/TAZ downstream genes confirm the activation of YAP/TAZ pathway in Group4_SCL cells. (**F**) Upper panel: JUN/TEAD4/YAP/TAZ ChIP-seq (GSE66083) data have stronger signal in Group4_SCL-specific ATAC-seq peaks while AR ChIP-seq (GSE52725) shows stronger signal in Group1_AR-specific ATAC-seq peaks. Lower panel: Venn diagrams showing the overlaps between the ChIP-seq peaks and Group1_AR- or Group4_SCL-specific ATAC-seq peaks.

#### Evidence from ATAC-seq and gene expression data

TFs in general work together and bind cooperatively in a context-specific manner to achieve specificity and execute their functions (*30*). Thus, to further investigate the regulation of Group4_SCL samples by AP-1, we investigated the other top TFs identified from chromatin accessibility profiles. We find that TEAD motifs are the second most enriched following AP-1 in the Group4_SCL-specific accessible peaks (fig. S17B) and they rank highly based on gain of chromatin accessibility and out-degree in Group4 samples (Fig. 5B, fig. S17A). The TEAD TFs are activated by YAP and TAZ transcriptional coactivators (*31*). In fact, motif analysis of the ChIP-seq peaks of YAP and TAZ has revealed that TEAD TFs are the main platform by which these proteins interact with DNA (*31*). TEAD, YAP and TAZ were reported to be associated with AP-1 genome-wide to transcriptionally co-operate in several contexts and jointly regulate the proliferation and motility in multiple cancers, including breast, colorectal and lung (*32*). In addition, TAZ (WWTR1) is among the top genes co-dependent with FOSL1 based on CRISPR (Avana) Public 20Q2 in DepMap (*33*) with a Pearson correlation of 0.33, further supporting the model in which FOSL1 functions together with YAP and TAZ. Based on ATAC-seq data and published literature, we hypothesized that YAP, TAZ, TEAD and AP-1 (FOSL1) may function together to promote the oncogenic growth of Group4_SCL tumors (Fig. 5C). This is supported by our observation that GSEA using the combined YAP and TAZ (YAP/TAZ) target signature defined by Wang *et al* (*34*) reveals strong enrichment (FDR < 0.001) in Group4_SCL compared to other samples (Fig. 5D). Group4_SCL also shows significantly higher expression of TAZ (fig. S17C) and qPCR analysis of representative YAP/TAZ target genes across the 24 samples shows their high expression in this group (Fig. 5E).

We find significant enrichment of overlaps between the HINT-ATAC predicted TEAD-bound peaks and AP-1-bound peaks in both DU145 and MSKPCa3, suggesting further that they may work together (p value < 0.001, Fisher’s exact test, fig. S17D, table S11). We use the master TF-bound peaks from other groups as a negative control and do not find significant enrichment of their overlap with TEAD-bound peaks (table S11). We also find significant overlap between the target genes of AP-1 and TEAD predicted in our regulatory networks (p value < 0.001, Fisher’s exact test, fig. S17E). Additionally, the predicted TEAD and AP-1 peaks are predominantly located at distal regions (fig. S18A), matching well with ChIP-seq results reported by others showing TEAD and AP-1 bind together at enhancer regions (*35, 36*).

#### Evidence from published ChIP-seq data

Furthermore, analysis of AP-1, TEAD, YAP and TAZ ChIP-seq peaks from multiple published cell lines (*35–38*) shows they exhibit large overlap with Group4_SCL ATAC-seq peaks but barely any with Group1_AR peaks (Fig. 5F and fig. S18B). Correspondingly, we also observe a strong enrichment of the ChIP-seq signal over Group4_SCL-specific ATAC-seq peaks compared to Group1_AR-specific peaks. As a negative control, the trend is opposite for AR ChIP-seq peaks, in which they show much larger overlap and signal enrichment at Group1_AR peaks compared to Group4_SCL as expected (*38, 39*). As a specific example, the ChIP-seq and ATAC-seq profiles illustrate open chromatin and binding of AP-1, YAP, TAZ and TEAD at enhancers of the representative YAP/TAZ target genes, *CYR61* and *AXL,* in Group4_SCL lines while the same loci are barely accessible in other groups (fig. S20 and S21).

### YAP/TAZ promote growth in Group4_SCL cells in a positive feedback loop with AP-1 and can be targeted by drugs

We used siRNA to knock down (KD) YAP and TAZ alone or together in DU145 cells. The knockdown efficiency was more than 80% based on qPCR and Western blot confirmation (Fig. 6A and 6B). Double knockdown of YAP and TAZ shows the strongest inhibition of canonical downstream targets (*CTGF, CYR61, AJUBA* and *ANKRD1*) as well as cell-cycle regulating genes *CCNA2* and *CCND1* (Fig. 6A and fig. S22A), which have been reported as YAP/TAZ targets in various model systems (*32, 36*). Similar results are observed in MSKPCa3 (fig. S22B and S22C). Importantly, upon YAP/TAZ double KD, we observe a significant decrease of cell growth in DU145 but not in the AR-dependent line 22Rv1 (Fig. 6C).

**Fig. 6.**
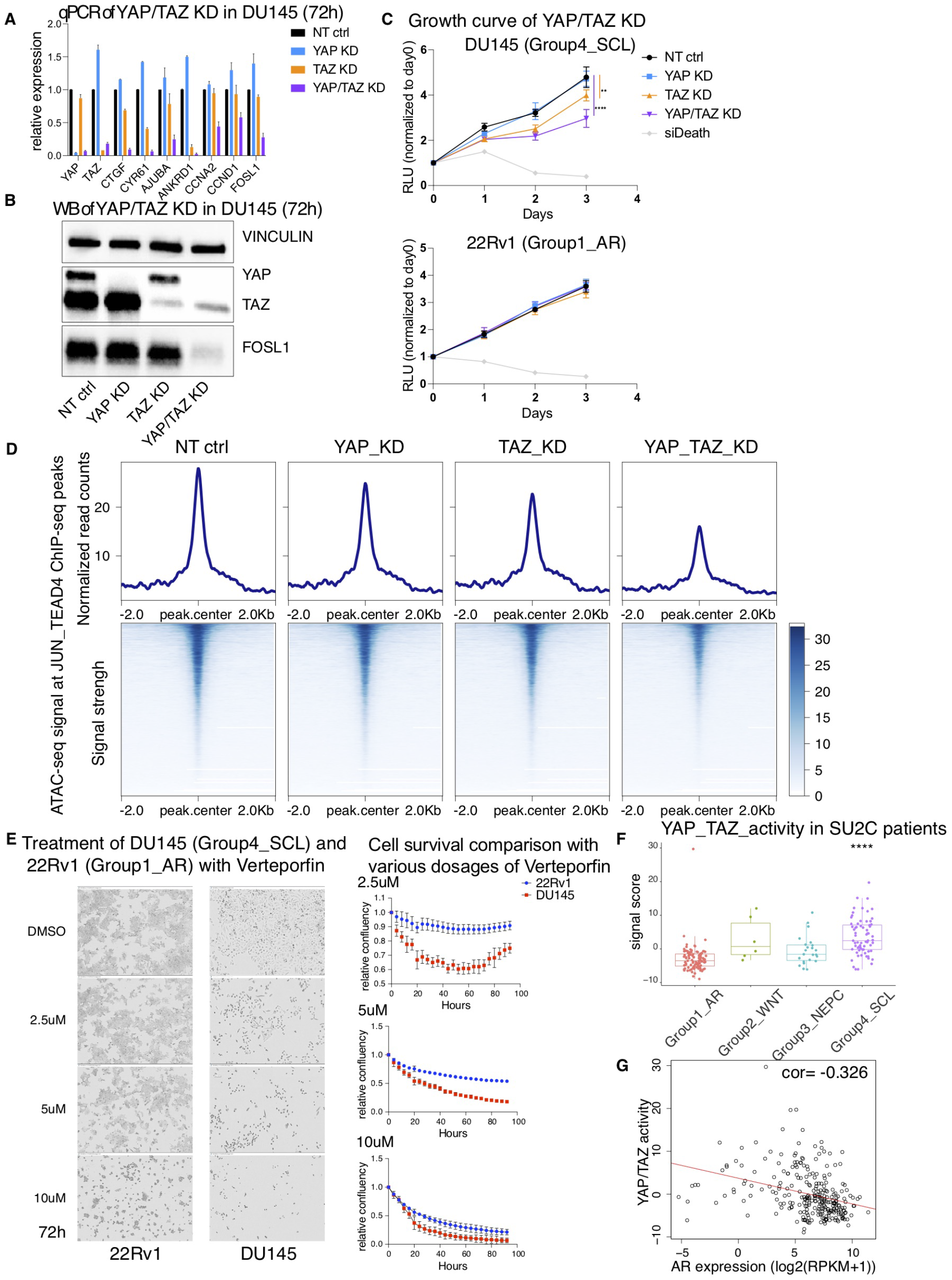
YAP/TAZ promote cell-growth in Group4_SCL cells in a positive feedback loop with AP-1 and can be targeted by drugs. (**A**) qPCR showing double knockdown (KD) of YAP/TAZ by siRNA in DU145 leads to the decreased expression of downstream target genes and *FOSL1.* Mean ± SD. Two-tailed unpaired t-test, n= 3. Non-targeting (NT) siRNA serves as the negative control. (**B**) Western blot (WB) confirming the KD efficiency and the decrease of FOSL1 expression 72h after transfection. (**C**) Cell growth curve of DU145 and 22Rv1 following siRNA KD. Mean ± SD. Two-tailed unpaired t test, n= 6. (**D**) The ATAC-seq signal is stronger over sites co-bound by AP-1 and TEAD, which is represented as the overlaps between JUN and TEAD4 ChIP-seq peaks (GSE66083) used in Fig 5F. (**E**) (Left) Representative of bright field images (10X) of DU145 or 22Rv1 treated with various dosages of Verteporfin or DMSO. (Right) Incucyte analysis showing that DU145 is more sensitive to Verteporfin than 22Rv1. At 72h timepoint, the maximum difference of relative confluency was observed at 5μM (56.4% in 22Rv1 and 23.5% in DU145). (**F**) YAP/TAZ activity (sum of z-scores) is significantly higher in Group4_SCL patients. The comparison was performed between the scores of each group to the overall signal scores across all patients. (**F**) YAP/TAZ activity is negatively correlated with AR expression across the 270 SU2C patients with cor=-0.326 and p-value < 0.001. ****p<0.0001.

Furthermore, we performed ATAC-seq on DU145 cells with KD of YAP and TAZ alone or together (Fig. 6D). We observe a significant decrease of chromatin accessibility at the sites cobound by AP-1 and TEAD upon double YAP/TAZ KD (Wilcoxon test, p-value < 0.0001) compared to non-targeting (NT) control. Moreover, we find that the TEAD and AP-1 motifs are the most enriched at the regions where chromatin accessibility is reduced upon YAP/TAZ double KD, further confirming the model in which YAP/TAZ co-operate with AP-1 and TEAD (fig. S22D).

Interestingly, YAP/TAZ double KD caused robust depletion of FOSL1, the predicted master TF, at RNA and protein levels in both DU145 and MSKPCa3 (Fig. 6A, 6B, fig. S22B and S22C). Moreover, in our regulatory networks for Group4_SCL samples, FOSL1 is predicted to be a target of FOS/JUN itself as well as TEAD which cooperates with FOSL1 in our proposed model. We also find that the Group4-specific FOSL1 enhancer is bound by TEAD, YAP, TAZ and FOS/JUN based on available ChIP-seq data (fig. S19). Together the results suggest that YAP, TAZ, TEAD and FOSL1 increase the expression of FOSL1 itself, forming a positive feedback loop to further open chromatin (fig. S22E).

Verteporfin is a benzoporphyrin derivative and a medication used as a photosensitizer approved by the FDA for the treatment of age-related macular degeneration. It has been widely reported to inhibit YAP/TAZ and cellular proliferation of multiple tumors (*40*). Consistent with the role of YAP/TAZ in Group4_SCL, we find DU145 cells are more sensitive to Verteporfin than 22Rv1 (Fig. 6E).

### YAP/TAZ pathway is enriched in Group4_SCL patients

Finally, we examined the YAP/TAZ activity in transcriptomic data from CRPC patients from both SU2C and WCM. YAP/TAZ pathway activity (sum of z-scores) is significantly higher in Group4_SCL patients relative to all samples (Fig. 6F, fig. S23A), with higher expression of YAP, TAZ and representative downstream genes (fig. S23B). We also observe a significant negative correlation between AR expression/AR pathway activity and YAP/TAZ pathway activity across all samples (Fig. 6G, fig. S23C).

## Discussion

We present a map of the chromatin accessibility and transcriptomic landscape of CRPC using a diverse biobank of organoids that recapitulate the genotypic and phenotypic heterogeneity of metastatic prostate cancer. We identify four subtypes of CRPC. By integrating ATAC-seq and RNA-seq data, we constructed sample-specific regulatory networks and predicted the master TFs for each subtype using a novel ranking algorithm. We further analyzed Group4_SCL because it constitutes the second most prevalent group in two large metastatic CRPC cohorts, exhibits significantly lower AR expression and AR transcriptional output, and is associated with worse overall survival under ARS1 treatment compared to Group1_AR.

Integrated analysis of ATAC-seq, RNA-seq and ChIP-seq data revealed a model in which YAP, TAZ, TEAD and AP-1 function together and drive oncogenic growth in Group4_SCL samples. Our model is validated by CRISPR and depletion studies using siRNA knockdown. This model enables us to go beyond the identification of master TFs that can be challenging to target by drugs (*41*) as it reveals therapeutic vulnerabilities by inhibition of the YAP/TAZ pathway.

Several prior studies support our conclusions. Liu *et al.* found that knocking down TAZ in DU145 (Group4_SCL from our study) decreased cell migration and metastasis, while overexpression of TAZ in RWPE (normal prostate cells) promoted cell migration, epithelial-mesenchymal transition and anchorage-independent growth (*42*). Overexpression of YAP was also reported to promote cell proliferation, invasion and castration-resistant growth in LNCaP (Group1_AR cells) and RWPE (*43*). YAP/TAZ activation has been found to be related to cell proliferation, therapy resistance, and metastasis in various types of tumors by extensive rewiring of the epigenome of differentiated cells, reprogramming them into stem-like cells and conferring lineage plasticity (*31, 32*). Our study is the first to show its important role in a specific subtype of CRPC.

Notably, the integrated use of ATAC-seq and RNA-seq data allowed us to identify the drivers of AR-neg/low CRPCs by pinpointing the master TFs. Previous studies based on RNA-seq data alone could not identify these drivers since GSEA identifies numerous biological processes that are enriched among Group4_SCL samples (fig. S3A), which complicates the efforts to find a predominant driver event.

We observe an enrichment of basal signature in Group4_SCL organoids (fig. S3C) and patient samples (fig. S23D). This basal signature has been observed in prostate cancer cell lines after depletion of TP53 and RB1 (*15, 18*) and is also observed in models of DNPC derived from AR knockout of luminal prostate cancer cells (*4*). This suggests that Group4_SCL tumors have acquired lineage plasticity similar to NEPC but are driven by different master TFs resulting in different phenotype. Because Group4_SCL are pathologically adenocarcinoma without neuroendocrine features, our study may guide the use of ARSIs in these cases. The mechanistic studies of lineage transformation upon depletion or over-expression of master TFs will be the focus of future studies. Overall, our study presents an approach to stratify CRPC patients into four subtypes using their transcriptomic signatures, which can potentially guide towards the most appropriate clinical decisions.

## Acknowledgements

We thank members of the Khurana and Chen laboratories for valuable critiques and discussions; the Genomic Core Facility at WCM for ATAC-seq and RNA-seq sequencing; the Englander Institute for Precision Medicine for WCM CRPC and patient and organoid data; the Center of Epigenetics Research at MSKCC for ATAC-seq; Integrated Genomics Operation core facility at MSKCC for MSK-IMPACT sequencing and Kenneth Chang at Cold Spring Harbor Laboratory for generously providing LentiV_sgRNA_Cas9_GFP (LgCG) vector and design of sgRNA sequences.

## Author contribution

F.T., Y.C. and E.K. conceived of and designed the project. F.T., Y.C. and E.K. wrote the manuscript with the help of all authors. S.W, C.J.L, W.D, D.G, W.A, A.G, M.F.B, P.C, H.I.S and Y.C. performed or supervised the derivation, maintenance and characterization of the ten new organoids. F.T., J.P. and C.H. performed sample processing and ATAC-seq library construction. F.T., C.K.W. and S.C. performed functional validation experiments. H.T. constructed shRNA knockdown vectors with the supervision from L.D. F.T. performed the majority of bioinformatic analyses. K.E performed whole exome sequencing data analysis for WCM organoids. R.H. provided bam files for WCM patient cohorts RNA-seq data. E.M.L. assisted with SU2C cohort analysis. A.P. assisted with the collection of organoids and cell lines SNV and CNV data. L.P. and H.B. provided WCM organoids. S.B., L.P., J.M.M, H.B., C.N.S. and A.S. collected and organized the WCM patient information and provided the sequencing data. Y.C. and E.K. supervised the study.

## Data and materials availability

All the data will be made available by submission to Gene Expression Omnibus.

## Supplementary Materials

Materials and Methods

Details of patient tumors for organoid derivation

Figs. S1 to S23

Tables S1 to S15

References

