## supplementary for "Chromatin accessibility profiles of castration-resistant prostate cancers reveal novel subtypes and therapeutic vulnerabilities"

### Materials and Methods

Organoid line derivation and culture

Metastatic prostate cancer biopsy tissues were collected with patient consent under Memorial Sloan-Kettering Cancer Center Institutional Review Board approval (MSKCC IRB 90-040, 06-107, or 16-865) prior to tissue acquisition.

MSKPCA 1-7 biopsy pieces were collected and minced with a sharp scalpel and digested for 0.5-2 hours at 37ºC rotating gently in a mixture of 5ml Advanced DMEM/F12 (ADMEM/F12) and 5mg/ml collagenase type II (Invitrogen). Dissociated cells were washed and seeded (*6*). MSKPCA 8-17 were derived from 68 biopsy samples. The details of the patient tumors characteristics and treatment history can be found in the "Details of patient tumors for organoid derivation" in the supplementary text. The biopsy pieces were collected and finely minced with a sharp scalpel in 50ul of human prostate organoid media(*44*). The minced tissue was then seeded directly into Matrigel.

All cells were seeded in growth factor reduced Matrigel (Corning) on 6 well low adherence plates (Greiner). If cells were successfully grown in Matrigel (3D) then cells were plated onto collagen (Rat tail 1 protein, Gibco) coated 10cm dishes (2D). Cultures were passaged when cells reached 80-90% confluency and split at a ratio of 1:3. The passage timeline varies for each cell line. Human prostate cancer organoids were dissociated using TrypLE (Invitrogen) for 5 minutes in 37ºC and/or with trituration using a specialized glass Pasteur pipet. Organoid lines were confirmed mycoplasma free by PCR testing.

Cell line source and culture

The LNCaP, 22Rv1, VCaP, DU145, PC3, C4-2 and H660 cell lines were obtained from the American Type Culture Collection (ATCC). All of these cell lines except for H660 were maintained in RPMI supplemented with 10% Fetal bovine serum (Omega), penicillin (100 U/ml), and streptomycin (100 ug/ml). H660 was grown in RPMI-1640 with 0.005mg/mL insulin (SCBT #sc-360248), 0.01 mg/mL transferrin (Sigma T5391-10MG), 30nM sodium selenite (Sigma S5261-10G), 10nM hydrocortisone (Sigma H0888-1G), 10nM B-estradiol (Sigma E2257-1MG), 2mL L-glutamine (Thermo Fisher 3505061), 5% FBS (Thermo Fisher 16000044), penicillin (100 U/ml), and streptomycin (100 ug/ml). Cell lines were confirmed mycoplasma free by PCR testing.

Immunoblot

Organoids and cell lines were lysed in RIPA buffer (Thermo Fisher 89901) supplemented with Halt Protease and Phosphatase Inhibitor Cocktail (Thermo Fisher, 78440). The total protein concentration of the supernatant was determined using the DC Protein Assay Kit, (Bio-Rad, 5000116). Each protein sample (20 µg/lane) was resolved on a 4-15% Mini-PROTEAN TGX Precast Protein Gel, (Bio-Rad, 4561086), transferred onto a nitrocellulose membrane, blocked, and incubated overnight at 4 °C with primary antibodies. Primary antibodies and their dilutions are listed in table S12. Following four washes with TBS-T, the blot was incubated with either horseradish peroxidase-conjugated anti-rabbit IgG (1: 10,000, Thermo Fisher, 32260) or horseradish peroxidase-conjugated anti-chicken IgY(H+L) (1:10,000, Thermo Fisher, 31401). After washing 4X protein bands were visualized by chemiluminescent detection using Immobilon Forte (Millipore WBLU0500).

ATAC-seq library preparation and data preprocessing

Culture conditions and sample processing were optimized for each organoid and cell line to reach >90% viability for sequencing. The libraries were generated using the OMNI-ATAC protocol(*45*), and the final library was purified with AMPure beads (Beckman Coulter, A63880). Library quality was verified using Agilent Bioanalyzer 2100 and Qubit (Thermo Fisher Scientific, Q32854). Samples were sequenced at WCM Genomic Core facility using Illumina HiSeq4000 with either 50 or 75bp paired-end reads and 40~50 million reads were generated for each sample, with two biological replicates for each.

The raw data were processed with the ENCODE ATAC-seq pipeline (https://github.com/ENCODE-DCC/atac-seq-pipeline). In short, the reads were trimmed, filtered and aligned against hg19 using Bowtie2 (*46*). PCR duplicates and reads mapped to the mitochondrial chromosome or repeated regions were removed. To correct for the Tn5 transposase insertion, mapped reads were shifted +4/-5. Peak calling was performed using MACS2 (*47*), with p value < 0.01 as cutoff. Reproducible peaks from two biological replicates were defined as peaks with Irreproducibility Discovery Rates (IDR) < 0.05. The distribution of inserted fragment lengths shows typical nucleosome banding patterns, and the TSS enrichment score (reads that are enriched around TSS against background) ranges between 8 and 35, suggesting the libraries have high quality and were able to capture the majority of regions of interests (fig. S2A to S2E, table S4).

RNA-seq library preparation and data preprocessing

RNA was isolated from organoids using Qiagen RNeasy kits. Library preparation and RNA sequencing was performed at MSKCC Genomics Core using Illumina HiSeq with 50 to 75bp paired-end reads and 20 to 50 million reads were generated per sample.

The sequencing data were mapped to the human reference genome (hg19) using STAR_2.4.2a (*48*) and annotated with GENCODE Release 10. Raw readcounts and transcripts per million (TPM) values were generated with featureCounts (*49*) v1.6.2 and RSEM (*50*) v1.3 respectively.

DNA Extraction and Submission of Samples for IMPACT Sequencing

80-90% confluent cells were collected from the Matrigel and spun down. TrypLE was then added for 5 minutes at 37$^{o}$C to dissociate cells from the Matrigel. When a clear separation of cells from Matrigel was detected, the Matrigel was aspirated out without compromising the cell pellet, and cells were then washed twice with 1 x PBS. Cells grown on collagen coated plates were collected by adding TrypLE to the plate to detach cells. All cells were counted using a Vicell automated cell counter. DNA was isolated from the organoids using a Qiagen Puregene Core Kit A according to the manufacturer's protocol for cultured cell extraction. Samples were submitted to MSKCC's Integrated Genomics Operation core facility where library preparation and IMPACT sequencing were performed.

For 7 patients (MSKPCa8~ MSKPCa14), DNA from the tumor biopsy and its matched organoids were clinically sequenced (table S2). As germline sequencing was unavailable in MSKPCa15-17, the somatic mutations for these organoids could only be evaluated using the unmatched method. This means that the normal DNA control was from a standard normal database, and therefore, the resulting mutations are likely the germline variants. The results are summarized in table S3.

Creation of ATAC-seq atlas and consensus clustering of ATAC-seq peaks
Reproducible peaks found in at least two of the samples were combined to create a chromatin accessibility atlas. The number of reads mapped to each peak was counted by featureCounts of Rsubread_1.32.4 (*49*) and a readcount matrix was created with 848,913 peaks as rows, and 53 samples as columns. The readcounts were normalized with variance-stabilized transform (vst) of DESeq2_1.22.2 (*51*). Bigwig files for the ATAC-seq peaks were generated with deepTools (*52*) 3.1.3. The peaks were normalized using the reciprocal of the sizefactor generated from DESeq2. Clustering analysis was performed with the top 1% variable peaks using ConsensusClusterPlus_1.46.0 (*53*). Consensus clustering was done using the k-means clustering algorithm for 1000 iterations with a resampling rate of 80%. The cumulative density function (CDF) of the consensus matrix (fig. S2G) for each k (number of clusters) was plotted (fig. S2H). The number of clusters was determined by the relative increase in area under the CDF curve, and the optimal k is the one that leads to no appreciable increase (fig. S2I). We chose four as the number of subtypes, according to the "elbow" point in the relative change in area under the cumulative consensus distribution function curve (fig. S2G and S2H).

To confirm the performance of the k-means clustering, hierarchical clustering, PCA and t-SNE analyses were performed with the top 1% variable peaks of the ATAC-seq peak atlas. The feature distribution of the ATAC-seq peaks was plotted with R package ChIPseeker (*54*) v1.18.0. There was no significant difference in the number of peaks across the four groups of samples (fig. S2I).

Differential gene expression and pathway analysis with RNA-seq data
Differential gene expression across the organoids and cell lines sequenced was calculated using DESeq2_1.22.2(*51*). FDR correction of 0.01 was imposed unless otherwise stated. To identify the pathways enriched in each group, the protein-coding genes were pre-ranked based on sign(log2FoldChange)*(-10log(pvalue)) when comparing one group of samples relative to others. GSEA was performed using JAVA GSEA_4.0.3 program (*55*) using pre-ranked gene list with 1000 permutations. Pathways used in the paper include the Broad Molecular Signatures Database gene sets (*55*) v7.1, h (hallmark gene sets) and c2 (curated gene sets), NEPC signature (*56*), YAP/TAZ pathway signature (*34*), TCGA AR signature (*57*), FGF signature (*4*), luminal signature (*15*), basal signature (*15*) and prostate cancer basal stem cell signature (*16*). The gene list in each of the signatures are listed in table S14. The pathway activity/signal score in organoids and cell lines in Fig.2G was calculated using GSVA_1.30.0(*58*) with mx.diff=F, min.sz=15, max.sz=500, method="gsva".

Analysis of the organoid whole-exome sequencing (WES)
FastQC (http://www.bioinformatics.babraham.ac.uk/projects/fastqc/) was run on raw reads to assess their quality. The output of FastQC gives several metrics, including average base quality of raw reads, sequence duplication of raw reads, and the k-mer enrichment along the length of raw reads. We kept these measures to confirm correct sequencing or de-multiplexing of the samples.

After initial QC, adapter sequences were trimmed using Trimmomatic (*59*). Short reads were then aligned to GRC37/hg19 reference using BWA. v6.2 with default parameters except –e 50 (to enable long indel detection). GATK v2.3.9 with default parameters was applied for a local realignment and base quality score recalibration of the mapped reads. This resulted into what we term a "clean" BAM file, ready for QC and variant calling. The alignment and analysis of the exome data was processed on Sun Grid Engine computing cluster using 8 CPU cores with 16Gb of memory.

Point mutations were detected using three separate approaches. The first approach used an in-house SNV caller, SNVseeqer (*60*) to determine which point mutations were found primarily in the tumor. Mutations found at positions reported in dbSNP (Build 137) were filtered out (*60*). As an additional filter, mutations were kept if they were located in coding sequences and caused an amino acid change determined by SNVseeqer. Indels were detected using GATK v2.3.9 somatic indel with default parameters. These mutations must be covered by at least 10 aligned reads in the tumor and a matched control. For the matched control, we sequenced blood from the patient that we derived the organoid from. Furthermore, the mutations were filtered by variant allele frequency where the mutation had to be present in the tumor with a variant allele frequency > 25% and present in the matched control with a variant allele frequency < 1%.

A second approach directly interrogated the tumor and matched control at positions reported in COSMIC. COSMIC is a database of somatic cancer mutations curated by the Sanger Institute (http://www.sanger.ac.uk). Positions reported more than ten times in the database were interrogated. At each of these positions, Samtools (*61*) interrogated the aligned reads in the tumor and matched control sample, and mutations with a variant allele frequency >5% in the tumor and <1% in the control were kept. These mutations must be covered by at least 30 aligned reads in both tumor and control samples. Lastly, mutations were filtered out if the SnpEff (*62*) annotation tool did not predict the mutation to cause an amino acid change (based on RefSeq gene annotation) based on the canonical transcript.

The third approach complements the second one and uses the same approach to look at tumor and control at positions where mutations are known *a priori* to be clinically relevant. These positions were generated through literature and database search and were placed in the clinically relevant category if a mutation in a specific genomic region was known to have sensitivity to an FDA approved drug(s). Samtools compared tumor and matched control samples and reported mutations with a variant allele frequency >5% in the tumor and < 1% in the control. Mutations must be covered by at least 30 aligned reads in both the tumor and control samples. Mutations predicted to cause an amino acid change by Annovar were kept.

For somatic copy number alterations, the number of aligned reads for capture regions in the HaloPlex Exome were calculated in both the tumor sample and matched control sample. Our rationale for taking this approach is that genomic regions that are aligned more frequently in the tumor sample relative to the control sample are indicative of copy number gains. Conversely, genomic regions that are aligned less frequently in the tumor sample relative to the control sample are indicative of copy number gains. Capture regions with a total coverage smaller than 100 reads in both the tumor sample and matched control sample were filtered out. For the remaining capture regions, read counts were normalized in both the tumor sample and the matched control sample by the total number of reads aligned in the tumor sample and the matched control sample respectively. Then the ratio of the normalized read counts in the tumor sample and the normalized read count in the control sample was calculated.

These capture regions were then ordered karyotypically and sorted by genomic coordinates to help segment our capture regions according to the log2 value of the ratio of normalized read counts of the tumor sample and control sample in a biologically meaningful way. The normalized ratios of these bins were segmented using the Circular Binary Segmentation algorithm implemented in the R package DNAcopy (*63*). The algorithm output segments such that every capture region found within these segments is represented by the same log2 value. This log2 value indicates whether the segment has DNA copy number gain (amplification) or DNA copy number loss (deletion). A negative log2 would suggest a segment was deleted and a positive value would suggest a segment was amplified. Segments with a log2 value > 0.5 were categorized as amplified and segments with a log2 value < -0.5 were categorized as deleted. We then took the segments called by the algorithm and with a custom script annotated these segments by RefSeq genes, whose transcription start and end sites overlap with the genomic coordinates assigned to these segments.

<u>Gene regulatory network inference</u>

To construct the regulatory network for each organoid and cell line, we first constructed peak-gene links (Fig.3A, step1). We used rlog normalized readcounts from DESeq2_1.22.2 (*51*) as input for RNA-seq data, then used chromVAR (*64*) to get ATAC-seq peaks of 500bp fixed width , followed by normalizing using vst. We overlapped peaks within +/- 0.5Mb of a gene's TSS and identified peak-gene links using FDR < 0.01 as the cutoff in a Pearson correlation test between normalized read counts of these peaks and the expression of the gene across all samples. In addition, we removed diffuse correlations where large genomic regions were highly co-accessible, with a similar protocol as the one used by the Corces RM *et al.* (*12*), in which we tiled the genome into 100-kbp windows, performed the vst normalization (*51*) of read-counts in peaks and computed the correlation between the 100-kbp window's total accessibility and the expression of the linked gene. If the correlation between a gene-peak pair was lower than the absolute correlation between the gene's expression and the 100-bp window where the peak summit resided, the peak-gene link was considered a diffused link and was removed for later analysis. In the end, peaks with summits less than 1500 bps away and targeting the same gene were merged together. In total, we identified 76,002 peak-gene links involving 8,981 genes, 4,752 of which are protein-coding.

In the second step (Fig.3A, step2), TF-peak connections were constructed by footprint analysis using HINT-ATAC (*21*) with the peak bigwig files, the IDR peaks, and curated motifs from CisBP (*65*) as input for each sample. We used peak-gene links and TF-peak links from the two steps to make TF-target gene edges (Fig.3A, step3), and a regulatory network was constructed for each of the samples (Fig.3A, step4).

<u>Identification of master transcription factor by TF_rank</u>

To identify the master TFs in each of the four groups of samples, we developed a scoring system to rank TFs (TF_rank) consisting of three metrics: 1) differential outdegree: *O_diff* ; 2) differential chromatin accessibility: *A_diff*; 3) Differential gene expression: *E_diff*.

$$O\_diff = \frac{\sum_{i=1}^{N1} log2\left(\frac{outdegree\_TF}{\sum outdegree}\right)_i}{N1} - \frac{\sum_{j=1}^{N2+N3+N4} log2\left(\frac{outdegree\_TF}{\sum outdegree}\right)_j}{N2+N3+N4}$$

$$E\_diff = -log10(pvalue) \times \frac{log2(FC)}{|log2(FC)|}$$

$$A\_diff = \frac{\sum_{i=1}^{N1} Zscore_i}{N1} - \frac{\sum_{j=1}^{N2+N3+N4} Zscore_j}{N2+N3+N4}$$

$$TF\_Rank = rank(O\_diff) + rank(E\_diff) + rank(A\_diff)$$

After inferring TF-gene regulatory networks for all samples, we identified the number of genes regulated by each TF (TF outdegree) in each sample. The outdegree of each TF in each sample was normalized by the sum of all TF outdegrees in that sample. We assume if a TF is important in one subtype of samples, it would regulate more genes in that subtype relative to other subtypes, as demonstrated with a higher *O_diff* (Out-degree difference). For each TF in each sample, we computed a z-score for the gain or loss of accessibility at its motifs relative to the average sample profile using chromVAR (*64*). A higher z-score suggests the TF is associated with more open chromatin, *A_diff* (Accessibility difference) was calculated as the average z-score difference between one group relative to others. We ranked TFs using differential expression, *E_diff* (Expression difference) as the 3rd criterion, based on the idea that TFs with higher relative expression are more important in that subtype of samples. This also allowed us to distinguish the contributions of various TFs from the same family with similar motif sequences. N1, N2, N3 and N4 are the number of samples in each of the four subtypes. Log2(FC) is the log2 value of fold change. Log2(FC) and pvalue are outputs from differential expression by DESeq2. We assigned ranks to the TFs independently based on the three metrics discussed above and added up the three ranks as the final TF_rank (Fig. 3D and fig. S8, table S7). We find AR has the highest TF_rank in Group1_AR subtype, providing support for our ranking criteria.

Patient sample preparation and sequencing data processing

For the Weill Cornell Medicine (WCM) patients, fresh tumor biopsy specimens were obtained prospectively through a clinical trial approved by the WCM Institutional Review Board (IRB). mRNA was extracted using RNeasy Mini Kit (QIAGEN) and Maxwell 16 LEV simplyRNA Tissue Kit. Total RNA integrity was checked using a 2100 Bioanalyzer (Agilent Technologies, Santa Clara, CA). RNA concentrations were measured using the NanoDrop system (Thermo Fisher Scientific, Inc., Waltham, MA). Preparation of RNA sample library and RNAseq were performed by the Genomics Core Laboratory at WCM. Specimens were prepared for RNA sequencing using TruSeq RNA Library Preparation Kit v2 as previously described (*6*). Each sample was then sequenced with the HiSeq 2500 to generate 2 × 75-bp paired-end reads. The sequencing data were processed in similar methods as the one used for organoid RNA-seq data. Batch effect correction was done with Combat from sva_3.30.1(*66*) package with regard to library prep method and biopsy sites.

The processed RNA-seq data, SNV and CNV calls for SU2C patient data were downloaded from cBioportal (www.cbioportal.org). The RNA-seq samples were batch corrected for biopsy site and institution with Combat from the sva package (*66*).

The AR score, NE score and YAP/TAZ activity were calculated by following a similar strategy as the TCGA study (*13*). Specifically, the pathway activity was calculated based on the sum of z-scores of the 20 AR genes (*57*), the genes up-regulated in NEPC patients (*56*), and the YAP/TAZ gene signature defined by Wang et al (*34*).

Assigning patients to subtypes using nearest template prediction

We derived the signature for each of the four subtypes of models as the genes with higher expression in one group relative to others using log2(FC)>1 and padj <0.01 from DESeq2_1.22.2 as the criteria. Genes with low expression or low variance in CRPC patient samples were filtered out, and the remaining signature genes were used in the following analysis. A template was constructed for each subtype by combining all the signature genes and assigning a value of 1 to the signature genes of that group, while 0 was assigned to the signature genes of other groups.

To match the CRPC patient samples to each subtype, we used the Nearest Template Prediction (NTP) algorithm to assign each patient to a molecular subtype using RNA-seq data. NTP involves computing the cosine distance (*d*) between each patient's RNA-seq data and each of the four templates and estimating the statistical significance based on a null distribution for *d* generated by randomly re-sampling the same number of genes from patient RNA-seq data 1000 times. The process was repeated four times for each patient to calculate the p-values for the four templates and false discovery rate (FDR) was used for multiple hypothesis correction Each patient was assigned to the group where it had the shortest, significant (FDR < 0.5) value for *d*.

Genomic association with outcomes in SU2C patients

Enrichment of genomic alterations in Fig.4D was performed with two-tailed, unpaired Fisher's exact tests in R_3.5.1. Comparisons of pathway activity or marker genes were performed with Wilcoxon ranked sum test relative to a reference group or all samples, as indicated in the figure legends. FDR was used for multiple hypothesis correction.

Overall survival analysis was performed for 52 of the 80 subjects available in cBioportal and they were assigned to Group1_AR or Group4_SCL in NTP analysis. Time on treatment analysis was evaluated for a subset of n = 41 patients among the 61 subjects (table S10) available in cBioportal (https://www.cbioportal.org). Kaplan–Meier analysis and p-values for survival/time on treatment of different groups of patients were generated from the log-rank statistic with R packages survminer_0.4.3 (https://rpkgs.datanovia.com/survminer/index.html) and survival_2.42-3 (https://github.com/therneau/survival).

Overlap subtype-specific ATAC-seq peaks with ChIP-seq signal

Subtype-specific ATAC-seq peaks used in Fig.5F and fig. S18B were defined as peaks with log2(FC)>2 and padj < 0.01 when comparing one group of samples to the rest. The published ChIP-seq peak bigwigs were downloaded from ChIP-Atlas (https://chip-atlas.org/peak_browser) with 50 as the threshold of significance. The bigwig files for the ATAC-seq peaks in each sample were generated with deepTools 3.1.3 (*52*) using the sizeFactor from DESeq2 (*51*) for normalization. The profiles were plotted with deepTools, with subtype-specific ATAC-seq peaks as bed file input and TF ChIP-seq bigwig files as the bigwig input. The signal tracks in fig. S19-S21 were generated with Integrated Genome Browser 9.1.4 (*67*).

Statistical testing of the overlap of predicted TF-bound peaks based on ATAC-seq data

Predicted TEAD, AP-1, ASCL1, NEUROD1, AR and TCF peaks were generated by HINT-ATAC when constructing regulatory networks for each sample. AP-1 peaks consist of JUN, JUNB, JUND, FOS, FOSB, FOSL1 and FOSL2 peaks. TCF peaks consist of TCF7, TCF7L1, TCF7L2 and LEF1 peaks. TEAD peaks consist of TEAD1, TEAD2 and TEAD4 peaks. In each sample, all the peaks predicted to be bound by master TFs (AR, ASCL1, NEUROD1, TCF, AP-1) were collected as union of peaks. In Group4_SCL samples, one tail Fisher's exact test was performed to determine whether AP-1-bound peaks have significant overlap with TEAD-bound peaks; in Group1_AR samples, the same test was performed for AR-bound peaks and TEAD-bound peaks. Similar test was performed for samples from the other two groups. Except for AP-1 in Group4_SCL samples, the other master TFs didn't have significant overlap with TEAD in other samples (Table S11).

Motif analysis of ATAC-seq data with Homer

Enriched motif analysis using Homer (*68*) (findMotifsGenme.pl) with 'size -given -mask' was applied to more accessible regions in Group4_SCL samples compared to others (log2FC>2, padj < 0.01), or in less accessible peaks upon YAP/TAZ double KD compared to NT ctrl (log2FC <0, padj < 0.05) with the non-differentially accessible peaks as background.

siRNA knockdown and cell proliferation assay

For the YAP/TAZ knockdown experiment, a pooled ON_TARGET plus siRNA targeting YAP1 or targeting WWTR1 (TAZ) (Dharmacon/GE Healthcreae) was transfected into DU145, 22Rv1 or MSKPCa3 using RNAiMax (Thermofisher). Non-targeting siRNA (Dharmacon/GE Healthcreae) was used as negative control while AllStars Hs Cell Death siRNA (Qiagen 1027299) was used as positive control. For cell proliferation assays, DU145 (3,000 cells/well) or 22Rv1 (10,000 cells/well) were plated in 96-well plates overnight, and were transfected with siRNA the following day after adhering to the plates. Viability was measured daily with the CellTiter-Glo (Promega) assay kit according to manufacturer's instructions. Viability data were expressed as mean ± standard deviation (SD). Raw luminescence units (RLU) were then normalized on a per plate basis to the mean values of the negative control. Multiple sample comparisons were calculated using pair-wise comparison with FDR adjusting for multiple hypothesis testing (in GraphPad Prism 8). Differences between values were considered statistically significant at a p value of less than 0.05.

RNA isolation and RT-qPCR

For DU145 and MSKPCa3 were transfected with siRNA in 6 well plates, TRIZOL (thermos fisher, 15-596-018) was used to extract RNA following the protocol 48h or 72h after siRNA transfection. For RT-qPCR, RNA was reverse transcribed using the qScript cDNA SuperMix (QuantaBio, 101414-106). Lightcycler 480 SYBR Green I (Roche, 04887352001) was used to run qPCR on a Roche real time PCR machine. The qPCR primers are listed in table S13.

CRISPR/Cas9-mediated knockout and cell competition assay

For CRISPR/Cas9 knockout of target genes, single guide RNA (sgRNA) sequences were designed by Dr. Kenneth Chang (Cold Spring Harbor Laboratory) focusing on the DNA binding domain of each gene. The sequences of sgRNAs are as follows:

sgRosa26: GAAGATGGGCGGGAGTCTTC
sgRPA3: GATGAATTGAGCTAGCATGC
sgFOSL1#1: CGGCCAAGTGCAGGAACCGG
sgFOSL1#2: CAAGTGCAGGAACCGGAGGA
sgFOSL1#3: GGAACTGACCGACTTCCTGC

These sgRNAs were cloned into LentiV_sgRNA_Cas9_GFP (LgCG), generously provided by Dr. Kenneth Chang. The correct sequence of each sgRNA was confirmed by Sanger sequencing. Lentiviral particles were generated by co-transfecting the lentiviral constructs with psPAX2 and pVSV.G into HEK293T cells using X-tremeGENE 9 (Roche). Viral supernatants were collected 24- to 48-hours post-transfection. Protein lysates were isolated 7 days after transduction in target cells for Western blotting.

For competition growth assay, cells were stably transduced for expression of GFP and guide RNAs targeting respective genes. FACS analysis was performed at regular intervals to assess percentage changes of GFP-positive cells over the course of the experiment. The percentage of GFP positive cells at each passage was normalized to the initial ratio post transfection.

<u>Verteporfin treatment of cell lines</u>

Verteporfin (Selleck) was dissolved in DMSO as required. DU145 (3,000 cells/well) or 22Rv1 (10,000 cells/well) were seeded in a 96 well plates on Day 0, then treated with various dosages of Verteporfin or DMSO on Day 1. Cultures were monitored using Incucyte live-cell imaging system (Essen Bioscience). The average confluency from 4 images per well was plotted in biological triplicate for each cell line and each condition using kinetic imaging confluence measurement at 4-hour time intervals.

**Details of patient tumors for organoid derivation**

MSK-PCa8 is derived from a patient who was diagnosed with Gleason 8 prostate adenocarcinoma and underwent initial radiation therapy to the prostate with androgen deprivation therapy (ADT). The patient later developed castration-resistant metastasis to the bone while on ADT and was treated with palliative radiation therapy to the bone followed by Sipuleucel-T. He subsequently had progression in the bone and a biopsy was taken from a bone lesion in the left ischium. In culture, the tumor cells formed organoids that showed poor differentiation with anaplastic features, recapitulating the biopsy specimen. Immunohistochemistry (IHC) showed that the organoids were androgen receptor (AR) positive. The established organoid line exhibits a doubling time of 3.5 days.

MSK-PCa9 is derived from a patient who was diagnosed with Gleason 9 (5+4) prostate adenocarcinoma. Initial scans revealed bone metastasis. The patient was administered ADT. The patient developed castration-resistant disease, new and worsening bone disease, and started Sipuleucel-T. He subsequently had progression in his pelvic lymph nodes and a biopsy was taken of the right pelvic lymph node. Pathology of the lymph node showed prostate adenocarcinoma. Organoid cultures recapitulated the adenocarcinoma features with microacinar structures, which matched the tumor sample. The established organoid line exhibits a doubling time of 4 days.

MSK-PCa10 was derived from a patient who presented with bony and hepatic metastases. The patient underwent a prostate biopsy that demonstrated adenocarcinoma of the prostate with a Gleason score of 8 (4+4). The patient was administered ADT. A biopsy taken of the liver showed prostate cancer with neuroendocrine differentiation that was positive for synaptophysin. Cultured organoids showed typical small cell carcinoma morphology with a high nucleus to cytoplasm ratio and stained positive for synaptophysin. The established organoid line exhibits a doubling time of 3 days.

MSK-PCa11 is derived from a patient who presented with extensive metastatic Gleason 8 (4+4) prostate adenocarcinoma with metastases involving the bone, lung, liver, lymph nodes, and sinus/brain. He was administered ADT followed by docetaxel. A biopsy of the liver was taken after completion of docetaxel. The organoids in culture showed poorly differentiated adenocarcinoma and were AR positive. The established organoid line exhibits a doubling time of 4 days.

MSK-PCa12 is derived from a patient who was diagnosed with Gleason 9 (4+5) prostate adenocarcinoma with bone metastases. The patient was treated with ADT and developed castration-resistant disease with rising prostate-specific antigen (PSA) and progression in the bone and lymph nodes. A biopsy was taken from a pelvic lymph node. The organoids were poorly differentiated with squamous features and displayed high AR and KRT5 staining. The established organoid line exhibits a doubling time of 1.5 days.

MSK-PCa13 is derived from a patient who was diagnosed with Gleason 8 (4+4) prostate adenocarcinoma that metastasized to the lymph nodes. The patient had clinical metastatic non-castration-resistant disease and was treatment-naïve when he underwent a biopsy of the retroperitoneal lymph nodes. The organoids displayed high grade adenocarcinoma with typical prostate lumen structure matching the parental tissue. The established organoid line exhibits a doubling time of 7 days.

MSK-PCa14 is derived from a patient who was diagnosed with Gleason 7 (4+3) prostate adenocarcinoma with local advancement to the lymph nodes. The patient was treated with monthly degarelix. He subsequently progressed and developed castration-resistant disease. A biopsy was taken of a pelvic lymph node. The organoids, positive for synaptophysin, exhibited typical small cell histology with high nuclear to cytoplasmic ratio, which represented the biopsy sample. The established organoid line exhibits a doubling time of 5 days.

MSK-PCa15 is derived from a patient who was diagnosed with Gleason 7 (4+3) prostate adenocarcinoma. The patient underwent initial brachytherapy followed by ADT, but soon developed castration-resistant prostate cancer and was administered abiraterone followed by enzalutamide. The cancer cells were isolated from ascites fluid and cultured in organoids. In culture, the tumor cells formed organoids that were poorly differentiated, had anaplastic features, were positive for KRT5 and had low AR expression. The biopsy cytology slide was absent. The established organoid line exhibits a doubling time of 5 days.

MSK-PCa16 is derived from a patient initially diagnosed with de novo metastatic prostate cancer to the bone. The patient started ADT and received palliative radiation to the spine and to the prostate. He then had castration-resistant progression to the liver and a biopsy was taken. The organoids displayed histology of adenocarcinoma. The biopsy showed poorly differentiated carcinoma with focal synaptophysin and chromogranin positivity. The established organoid line exhibits a doubling time of 3 days.

MSK-PCa17 is derived from a patient with Gleason 9 (4+5) prostate adenocarcinoma. The patient underwent initial prostatectomy with positive surgical margins followed by ADT and salvage radiation therapy to the prostate bed. He then developed a biochemical relapse and non-castration-resistant metastatic disease to the bone. The patient was then treated with nilutamide, but developed castration-resistant disease and was administered abiraterone+cabazitaxel. A biopsy of the liver was taken while the patient was still on abiraterone but off cabazitaxel. The organoid showed adenocarcinoma. The biopsy tissue showed poorly differentiated adenocarcinoma. The established organoid line exhibits a doubling time of 6 days.

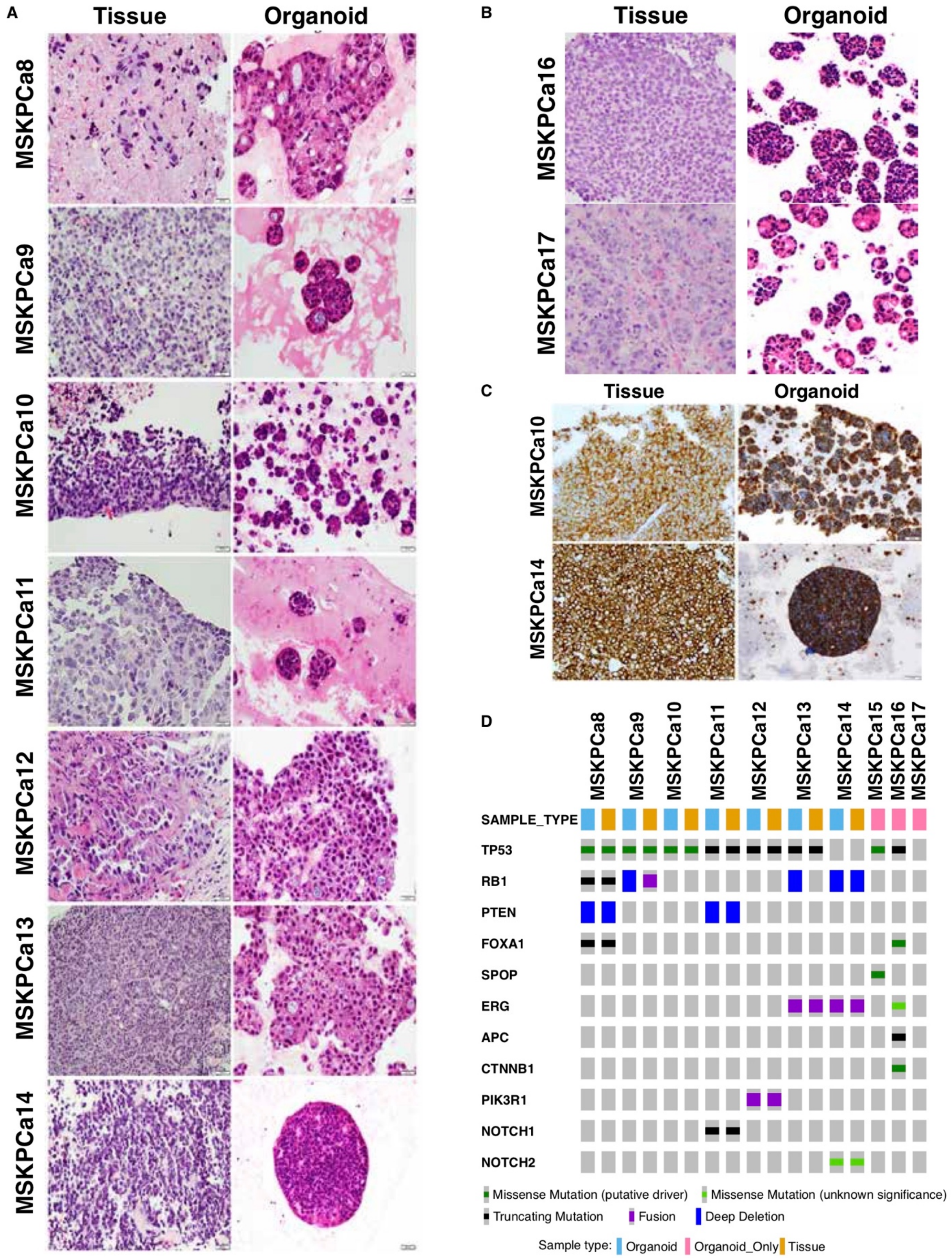

**Fig. S1 Characterization of organoids at genomic and pathological levels.**

(**A**) H&E staining for the tissue and organoid of MSKPCa8 ~ MSKPCa14. (B) H&E staining for the tissue and organoids of MSKPCa16 and MSKPCa17. (**C**) IHC for SYP are shown for MSKPCa10 and MSKPCa14 tissue and organoids. (**D**) Oncoprint showing the genomic alterations of selected genes of the ten organoids and seven matched tumor samples.

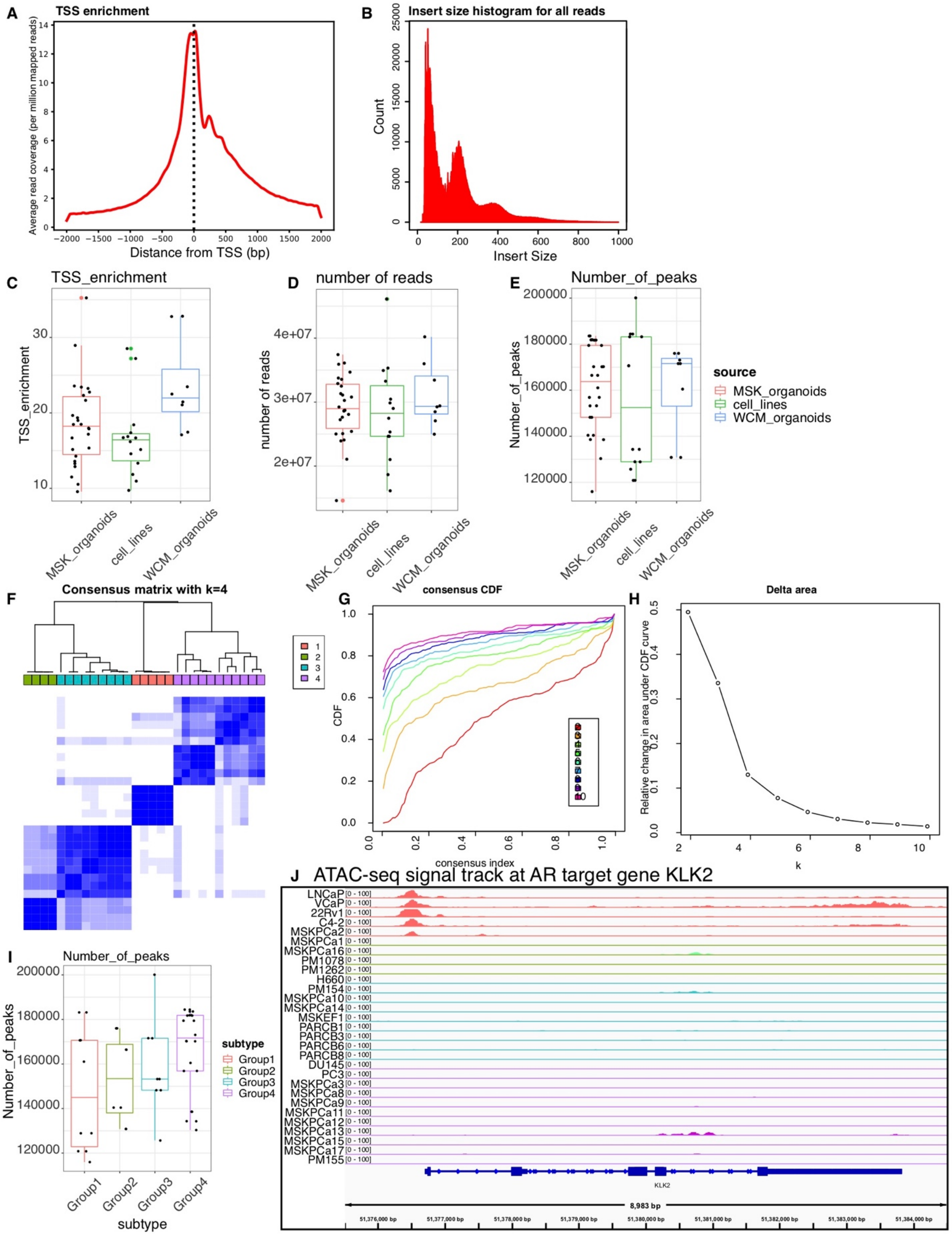

**Fig. S2 ATAC-seq data quality control and sample clustering.** (**A**) ATAC-seq signal enrichment around the transcription start site (TSS) in MSKPCa17. (**B**) The inserted fragment size distribution of MSKPCa17. (**C**) The distribution of TSS enrichment score distribution across all samples. (**D**) The distribution of the number of reads after alignment and filtering across all samples. (**E**) The distribution of the number of peaks across all samples. (**F**) Heatmap displays the consensus matrix of k-means consensus clustering on the basis of the ATAC-seq peaks (4 clusters). (**G**) Consensus empirical cumulative distribution function (CDF) of all given cluster numbers. (**H**) Relative change in area under CDF curve with the number of clusters suggests the optimal cluster number is 4. (**I**) There is no significant difference in the number of ATAC-seq peaks across the four groups of samples (Wilcoxon, two-sided test). (**J**) Normalized ATAC-seq sequencing signal tracks of 29 samples at the KLK2 locus.

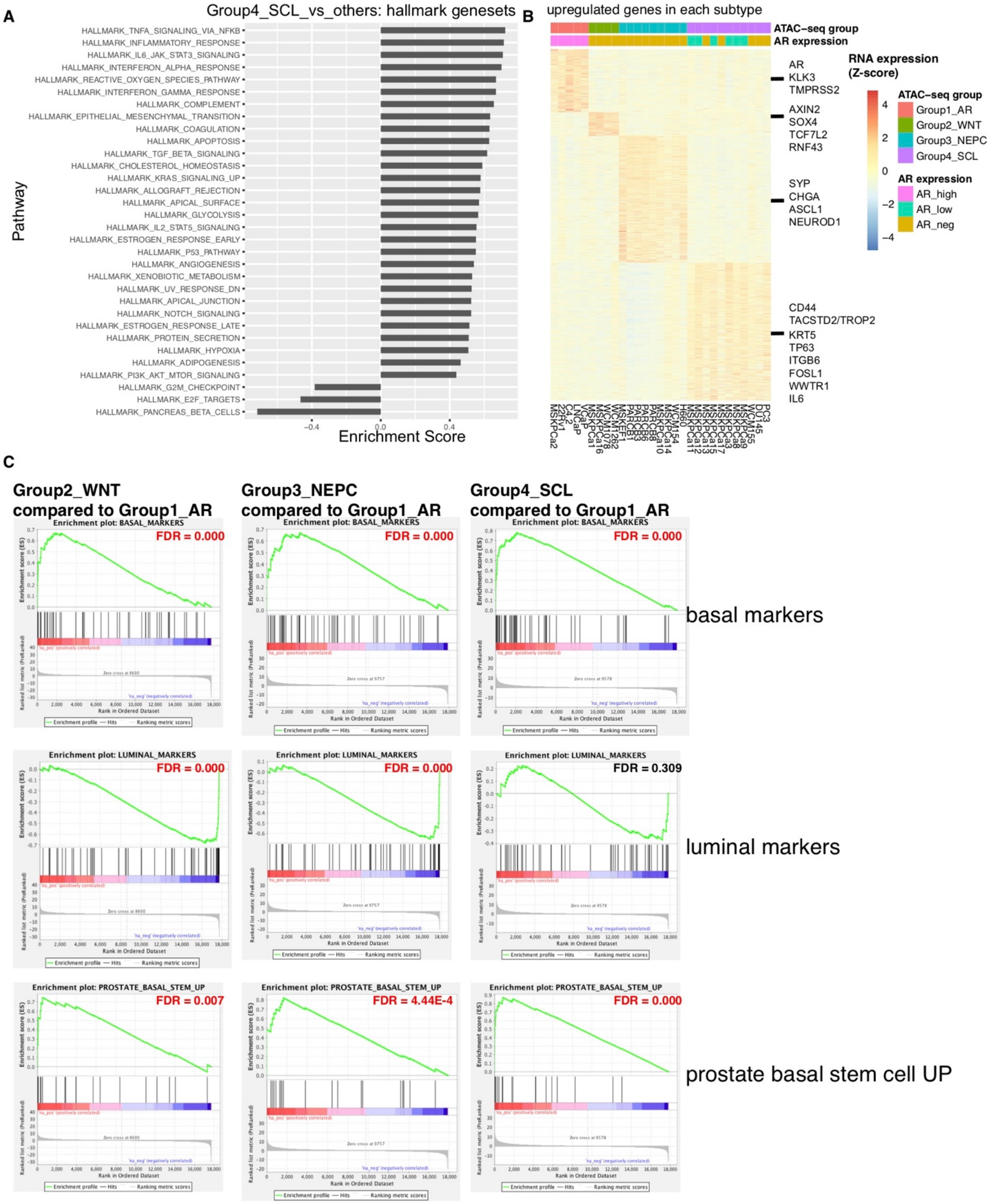

**Fig. S3. Pathway enrichment and differential gene analysis with the organoid and cell line RNA-seq data.** (**A**) The Enrichment Score of all the hallmark pathways that are significantly enriched or depleted in Group4_SCL samples compared to the others. (**B**) Heatmap showing the RNA expression of the upregulated genes in each of the four groups (log2FC >1, padj < 0.05) when compared between one group and all others. (**C**) GSEAs show the enrichment or depletion of basal signature, luminal signature and prostate basal stem cell signature of the non-AR-dependent subtypes relative to Group1_AR samples. FDR values < 0.05 are labeled red.

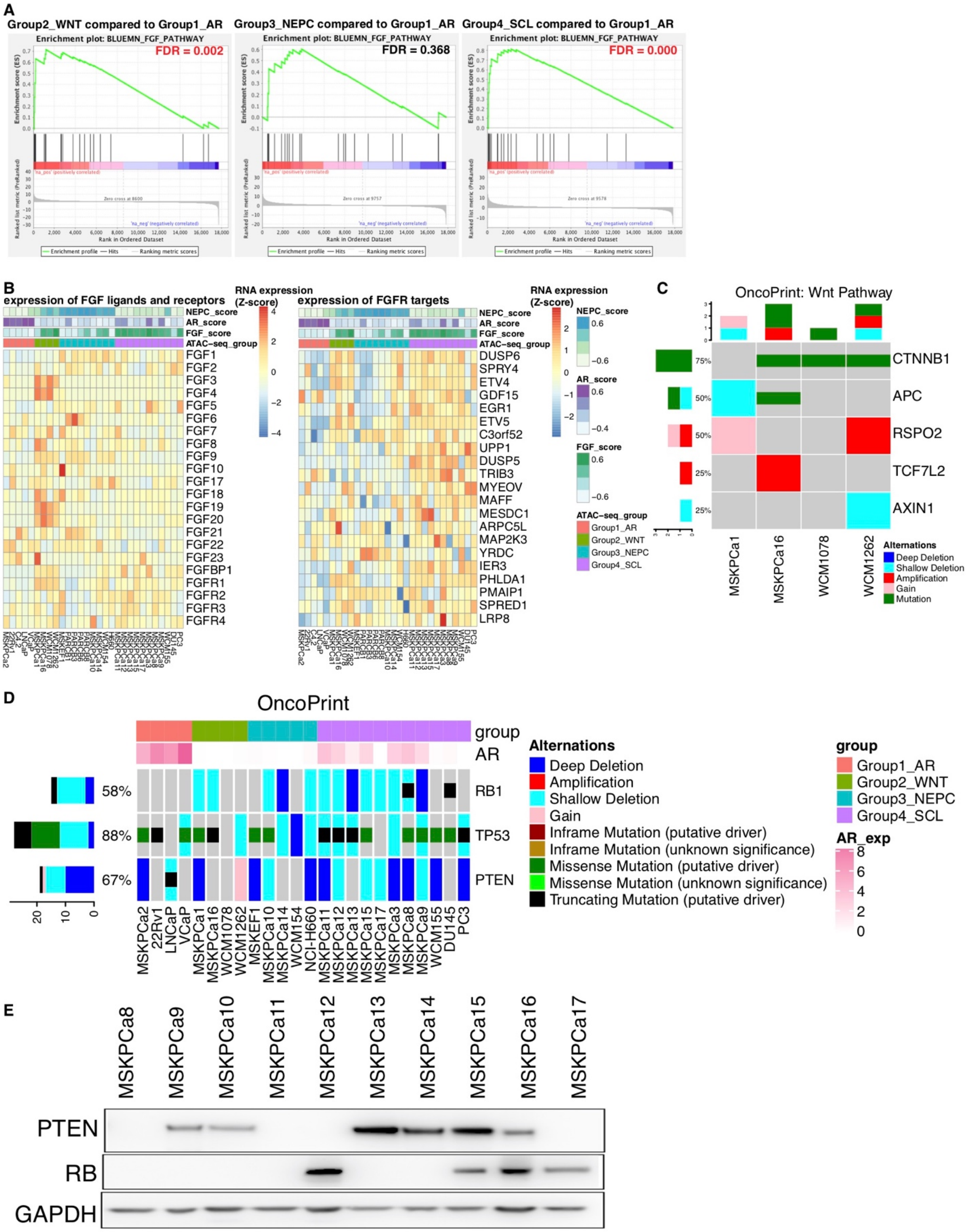

**Fig. S4. RNA-seq, DNA-seq and immunoblot of selective genes and pathways in organoids and cell lines.** (**A**) Both Group2_WNT and Group4_SCL samples have a significant enrichment of FGF signaling compared to Group1_AR. (**B**) Heatmaps showing the relative expression of FGF ligands, receptors (left) and downstream genes (right) across all samples. (**C**) The genomic alterations of Wnt pathway components are identified in all four samples in Group2_WNT. (**D**) Oncoprint showing the frequent alterations of RB1, TP53 and PTEN across the 24 samples from DNA sequencing data. (**E**) Western blot showing the expression of RB1 and PTEN across all ten newly derived organoids as stated above.

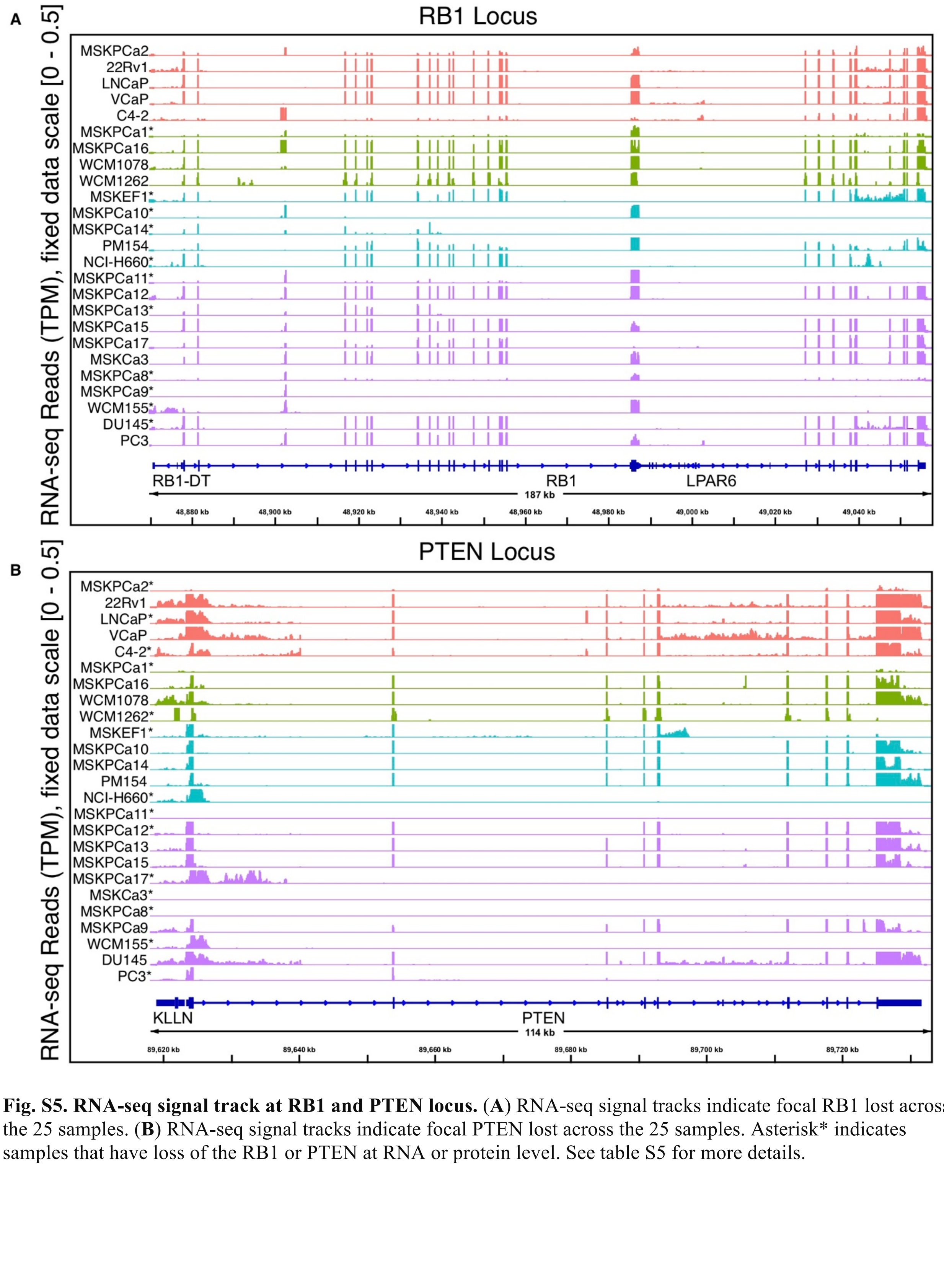

**Fig. S5. RNA-seq signal track at RB1 and PTEN locus.** (**A**) RNA-seq signal tracks indicate focal RB1 lost across the 25 samples. (**B**) RNA-seq signal tracks indicate focal PTEN lost across the 25 samples. Asterisk* indicates samples that have loss of the RB1 or PTEN at RNA or protein level. See table S5 for more details.

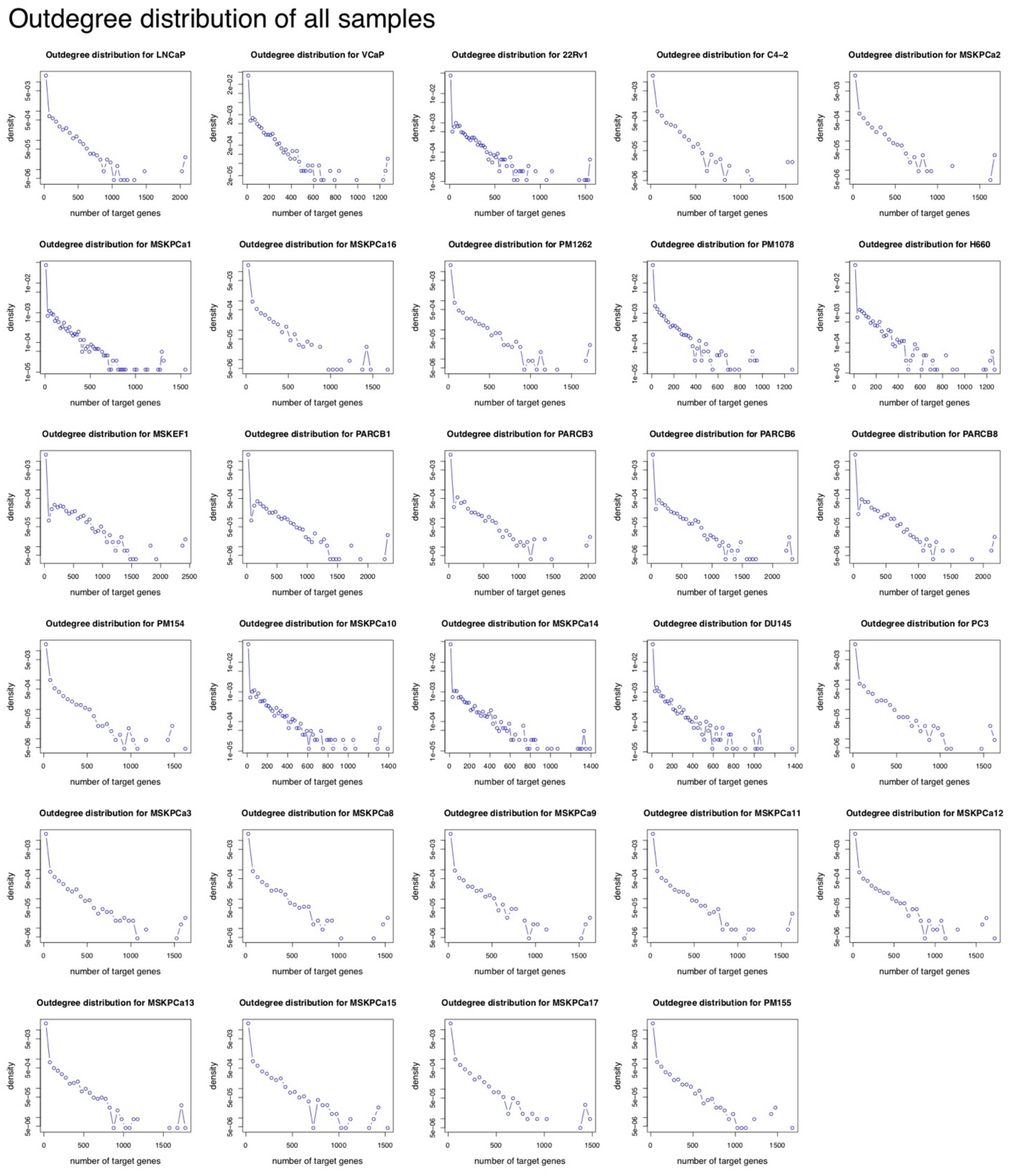

**Fig. S6. The out-degree distribution of the regulatory network in each sample.**

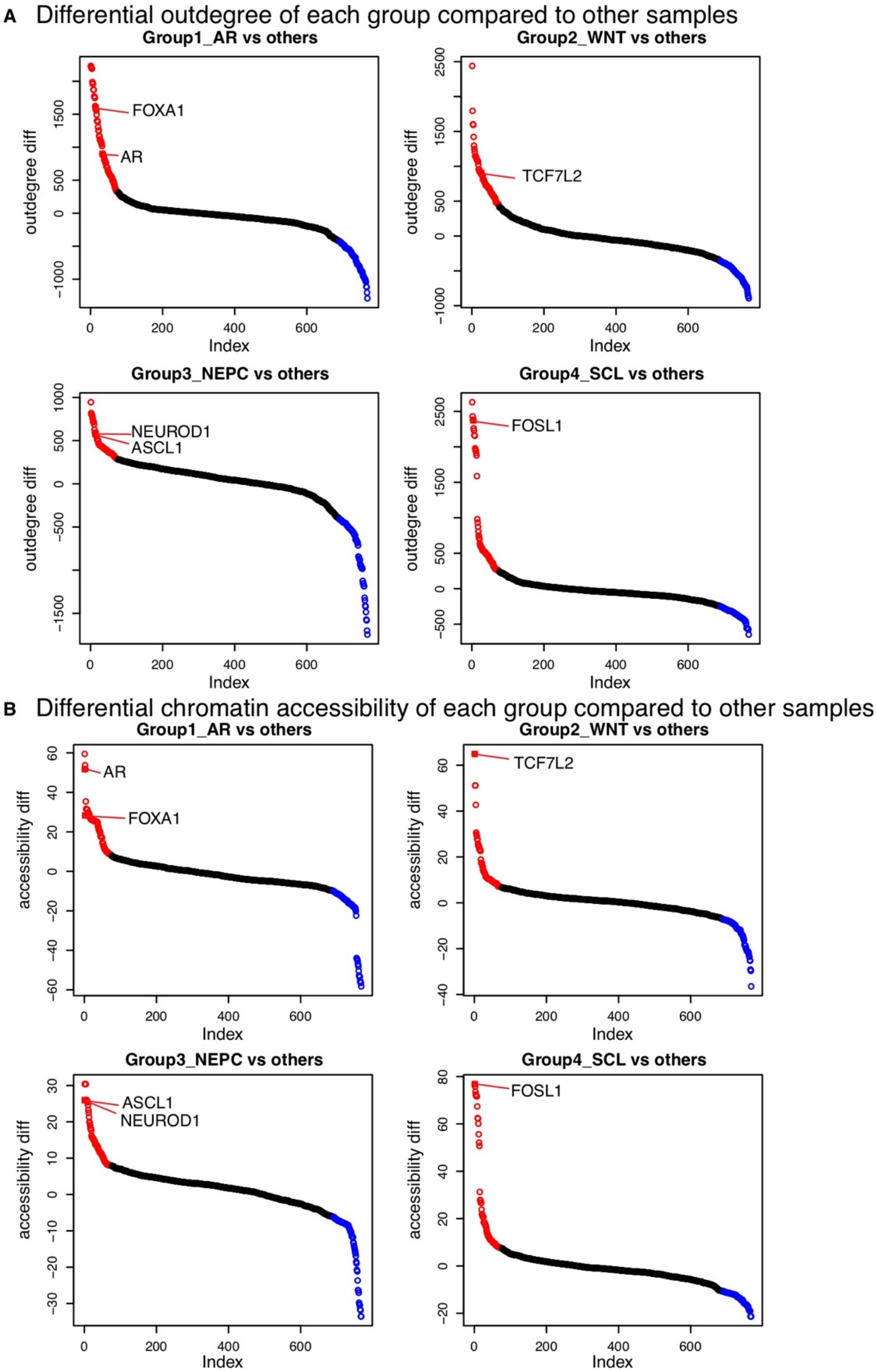

**Fig. S7. Differential outdegree and differential accessibility for all TFs in each of the four subtypes.** (**A**) Differential out-degree of TFs in each of the four subtypes compared to others. (**B**) Differential chromatin accessibility (zscore) in of TFs in each of the four subtypes compared to others. In (**A**) and (**B**) Each dot represents an individual TF. The top 10% and bottom 10% ranked TFs are labeled as red and blue circles respectively. The predicted final master TFs in each group are highlighted and labeled in red.

### 3D plot of the three metrics of all TFs in each subtype

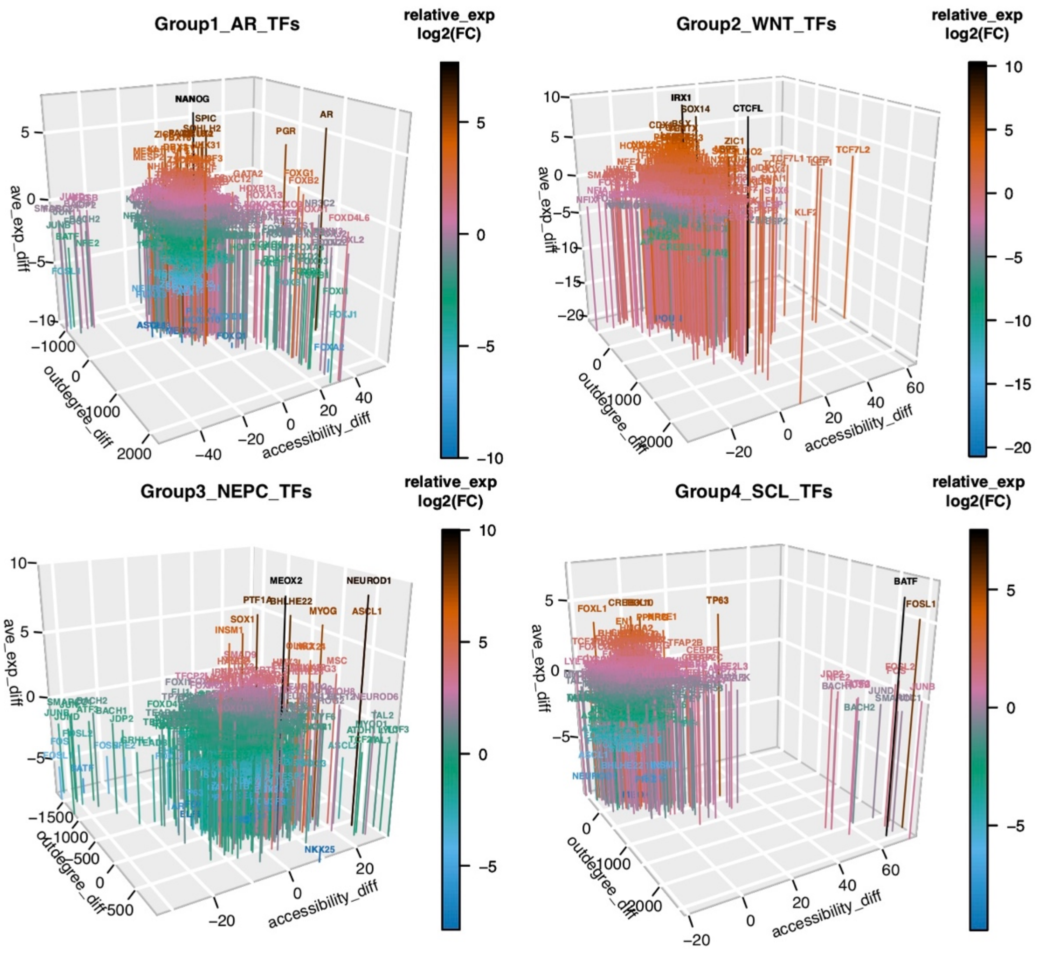

**Fig. S8. 3D plot of the three metrics of all TFs in each subtype.** The 3 metrics include differential chromatin accessibility (accessibility_diff), differential out-degree (outdegree_diff), and differential gene expression (ave_exp_diff) of TFs in each of the four subtypes compared to others.

Pairwise comparison between non-AR dependent samples to AR-dependent samples
(TFs with the top/bottom 20% differential zscore and expression were labeled)

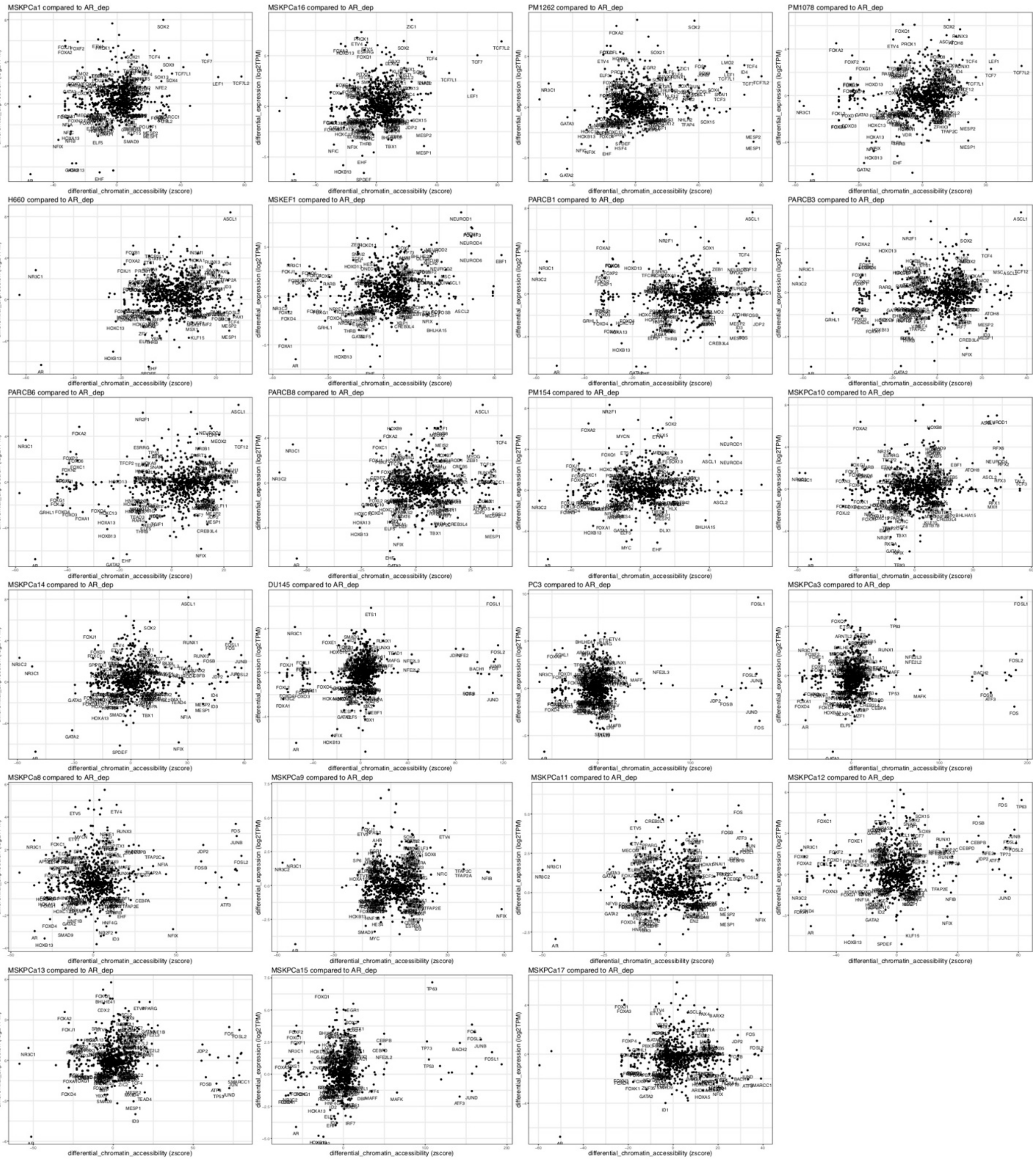

**Fig. S9**. **Pairwise comparison between each of the non-AR dependent samples and the average values of the AR-dependent samples.** The differential expression was plotted on the y-axis and differential chromatin accessibility (zscore from chromVAR) was plotted on the x-axis. TFs with the top/bottom 20% differential zscore and expression were labeled.

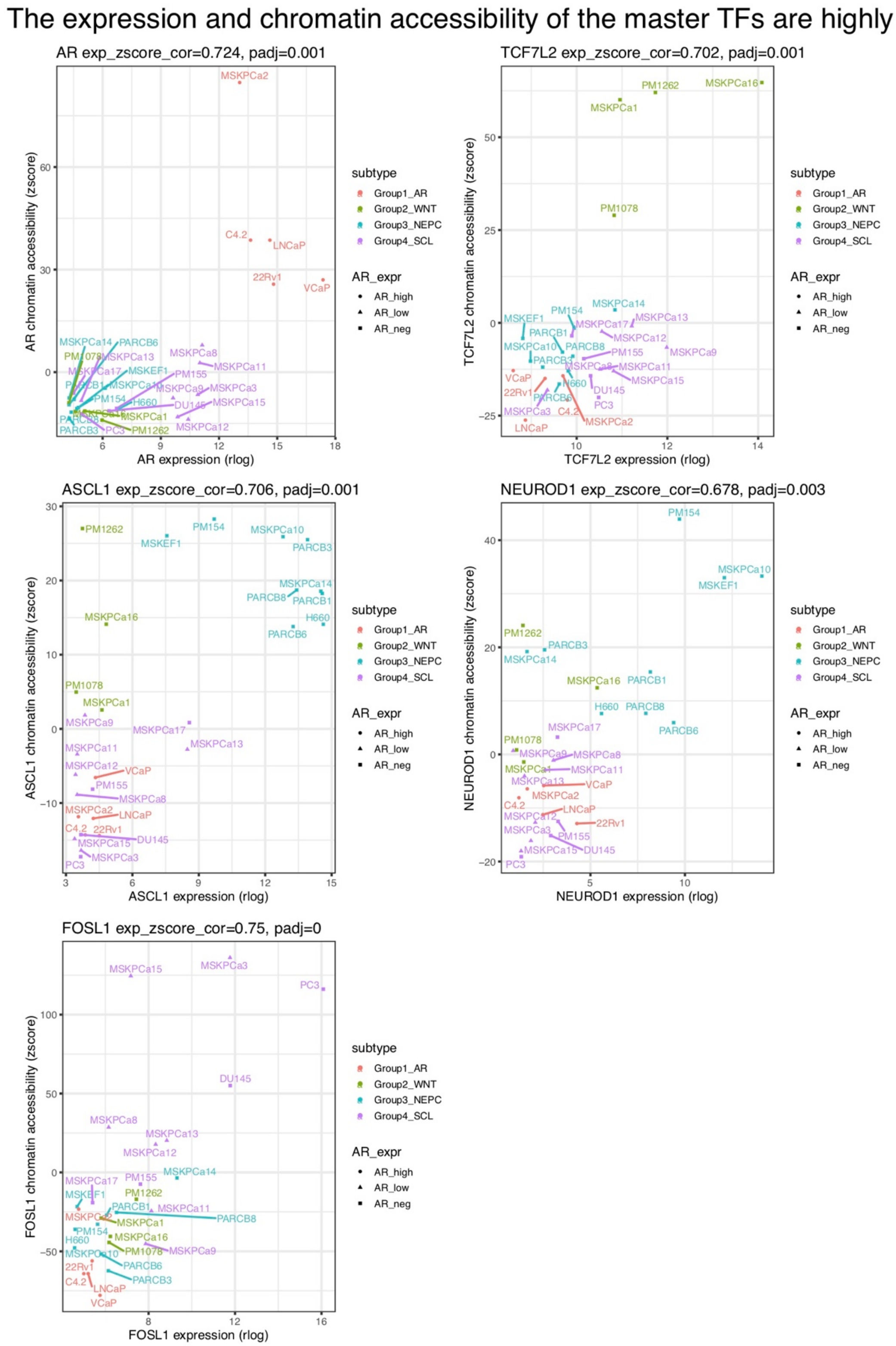

**Fig. S10. The expression and chromatin accessibility of the master TFs are highly correlated.** Chromatin accessibility of each TF (zscore from chromVAR) was plotted on the y-axis, and the expression (rlog from DESeq2) was plotted on the x-axis across all samples. FDR was used for multiple hypothesis correction.

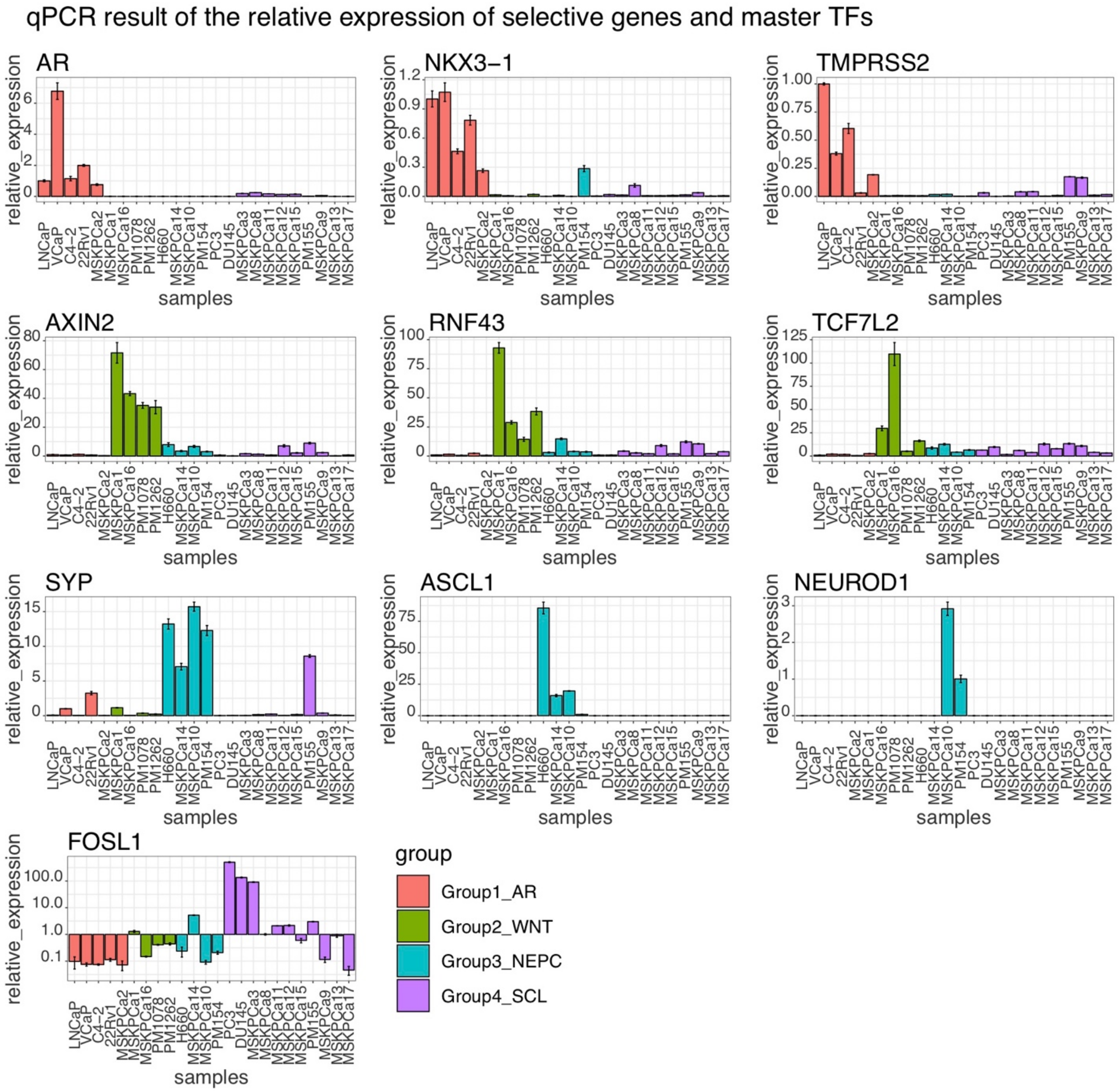

**Fig. S11. qPCR results showing the relative expression of representative marker genes and the master transcription factors of the four subtypes across all samples.**

### Dotplot showing distances of all SU2C samples to the four signatures in NTP analysis

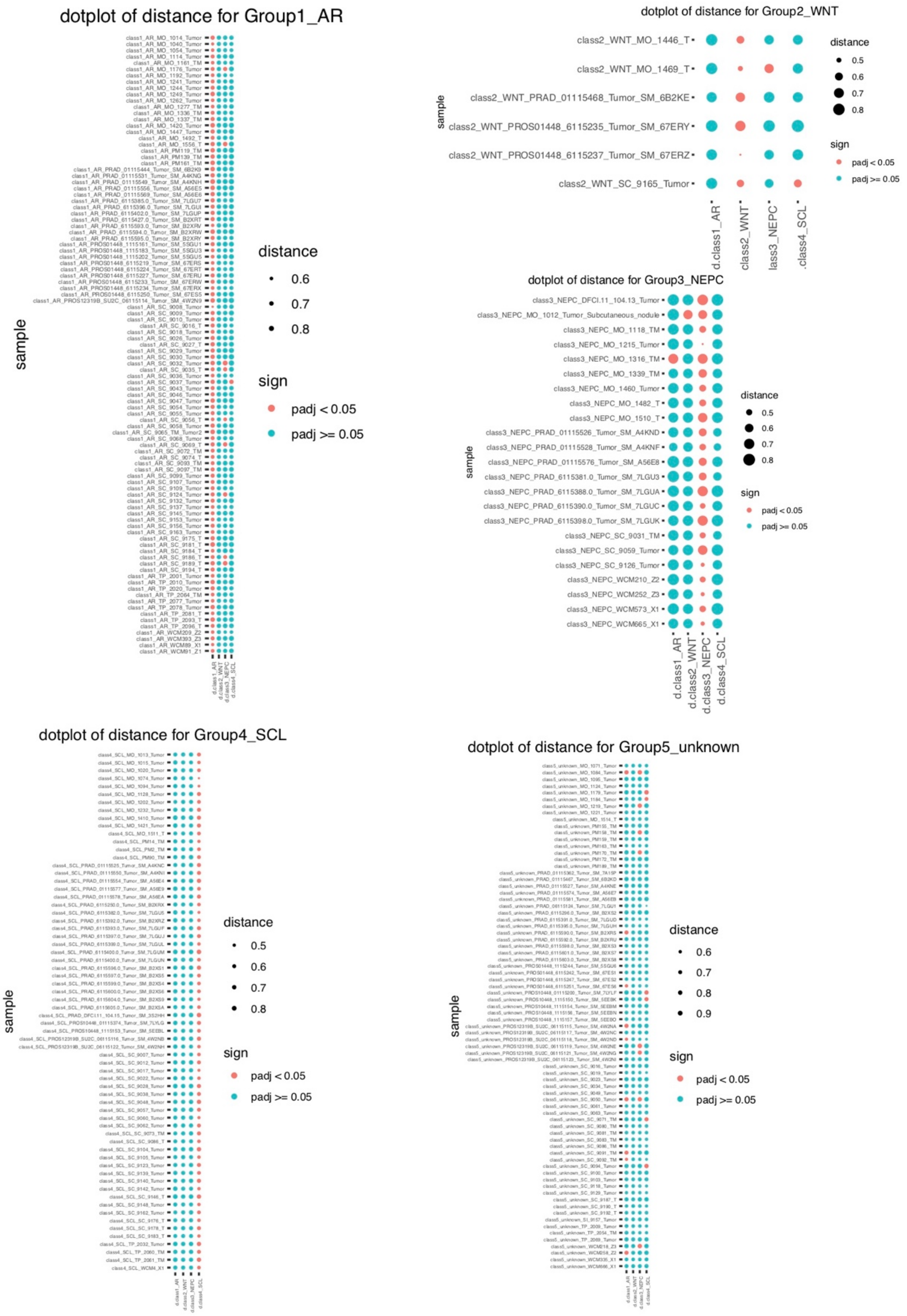

**Fig. S12. Dotplot showing distances of 270 SU2C samples to the four signatures in NTP analysis.** The radius of the dot indicates the cosine distance and the color indicates significance.

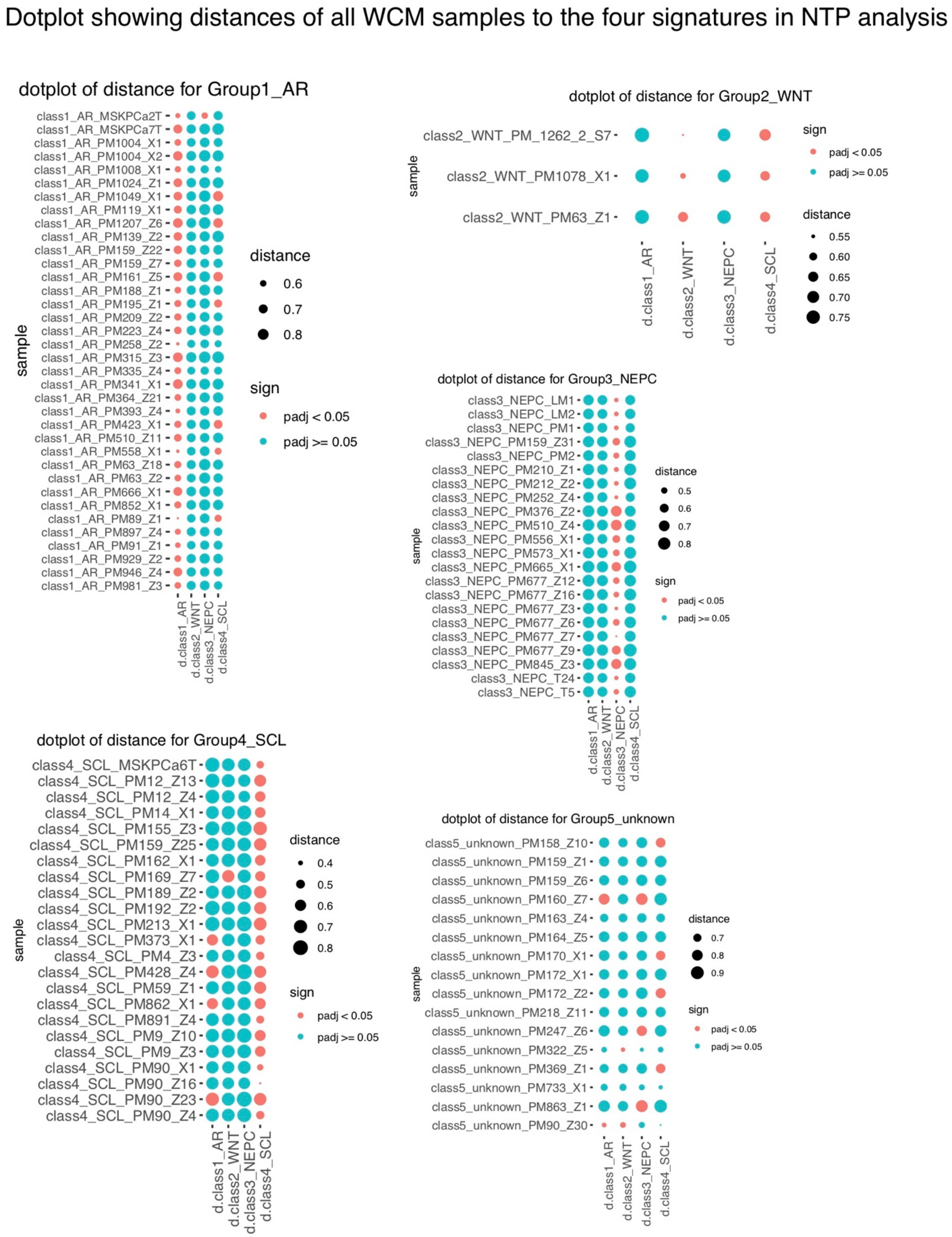

**Fig. S13. Dotplot showing distances of 100 WCM samples to the four signatures in NTP analysis.** The radius of the dot indicates the cosine distance and the color indicates significance.

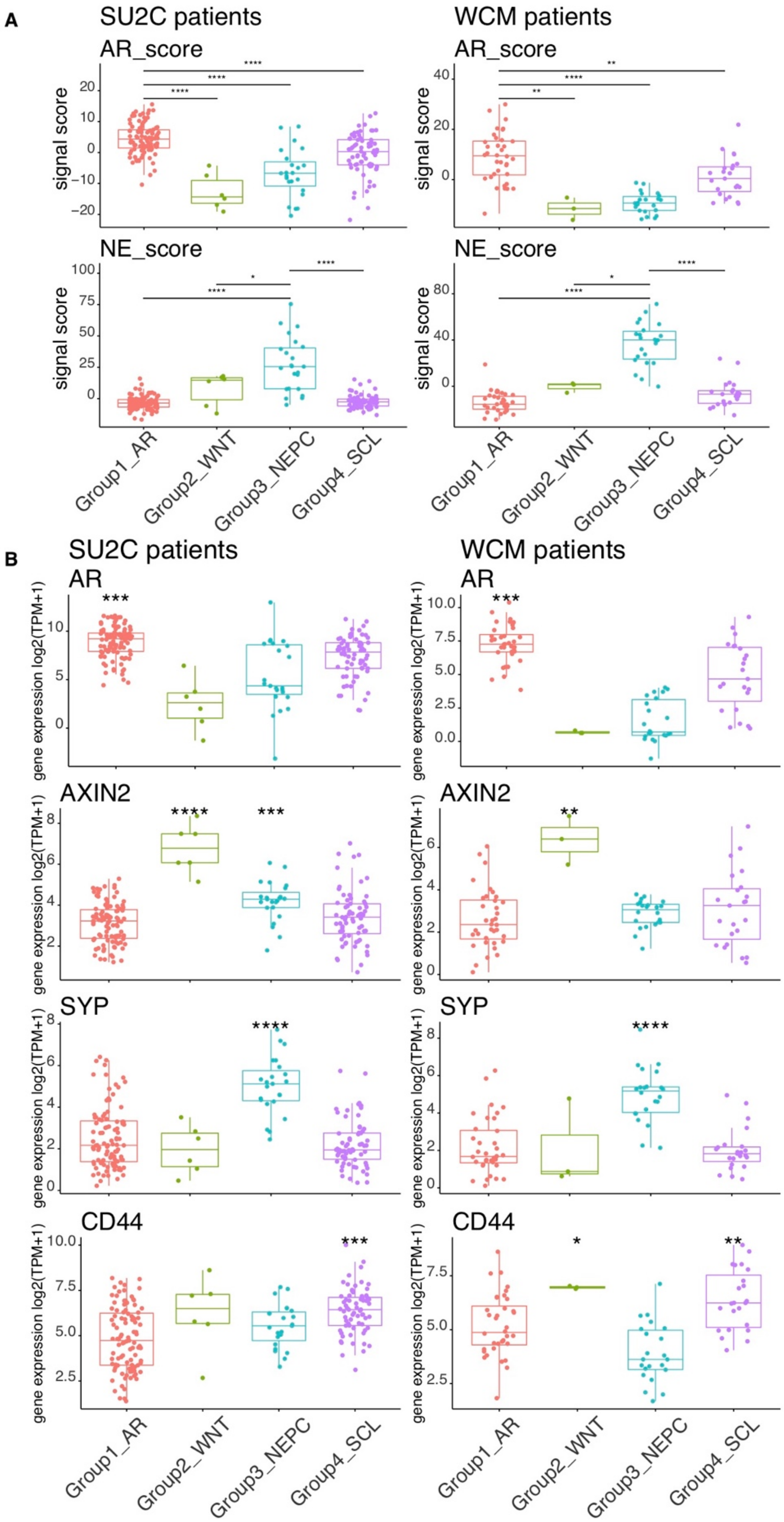

**Fig. S14. Comparison of pathway activity and marker genes among four subtypes of patients.** (**A**) Group1_AR and Group3_NEPC patients have significantly higher AR_scores and NE_scores respectively compared to other groups of patients. (**B**) The expression of the four representative marker genes for each group (AR, AXIN2, SYP,

CD44) are significantly higher in their corresponding subtype of patients compared to the overall expression of all patients. Adjusted p-values are calculated with one-sided Wilcoxon test and corrected with FDR. *p<0.05, **p<0.01, ***p<0.001, ****p<0.0001.

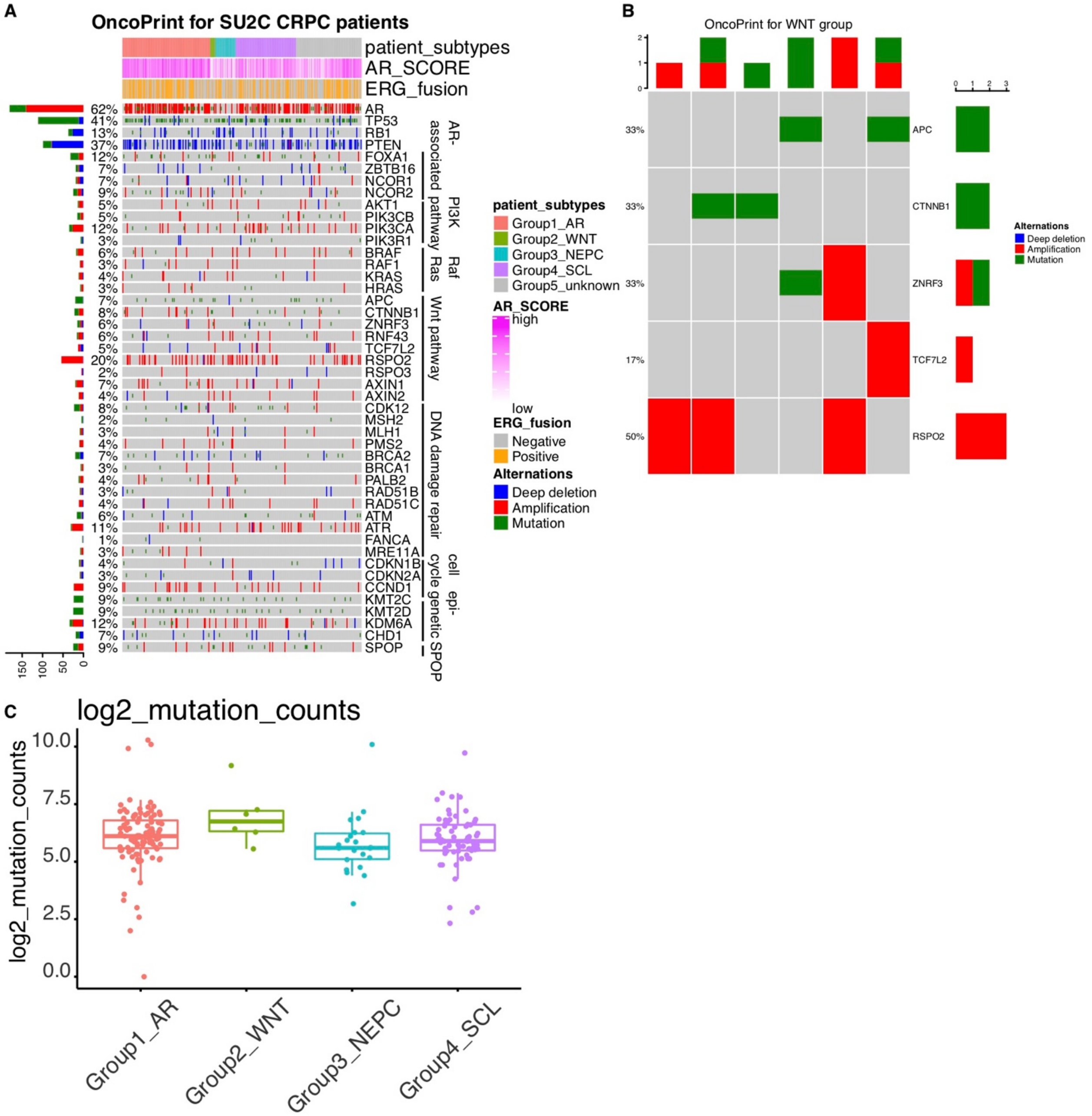

**Fig. S15. Genomic alterations in SU2C patients.** (**A**) Oncoprint showing the genomic alterations of selective genes across the 270 SU2C patients. (**B**) Oncoprint showing the alterations of Wnt pathway components across all Group2_WNT patients. (**C**) There's no significant difference in the number of mutations across all four patient subtypes.

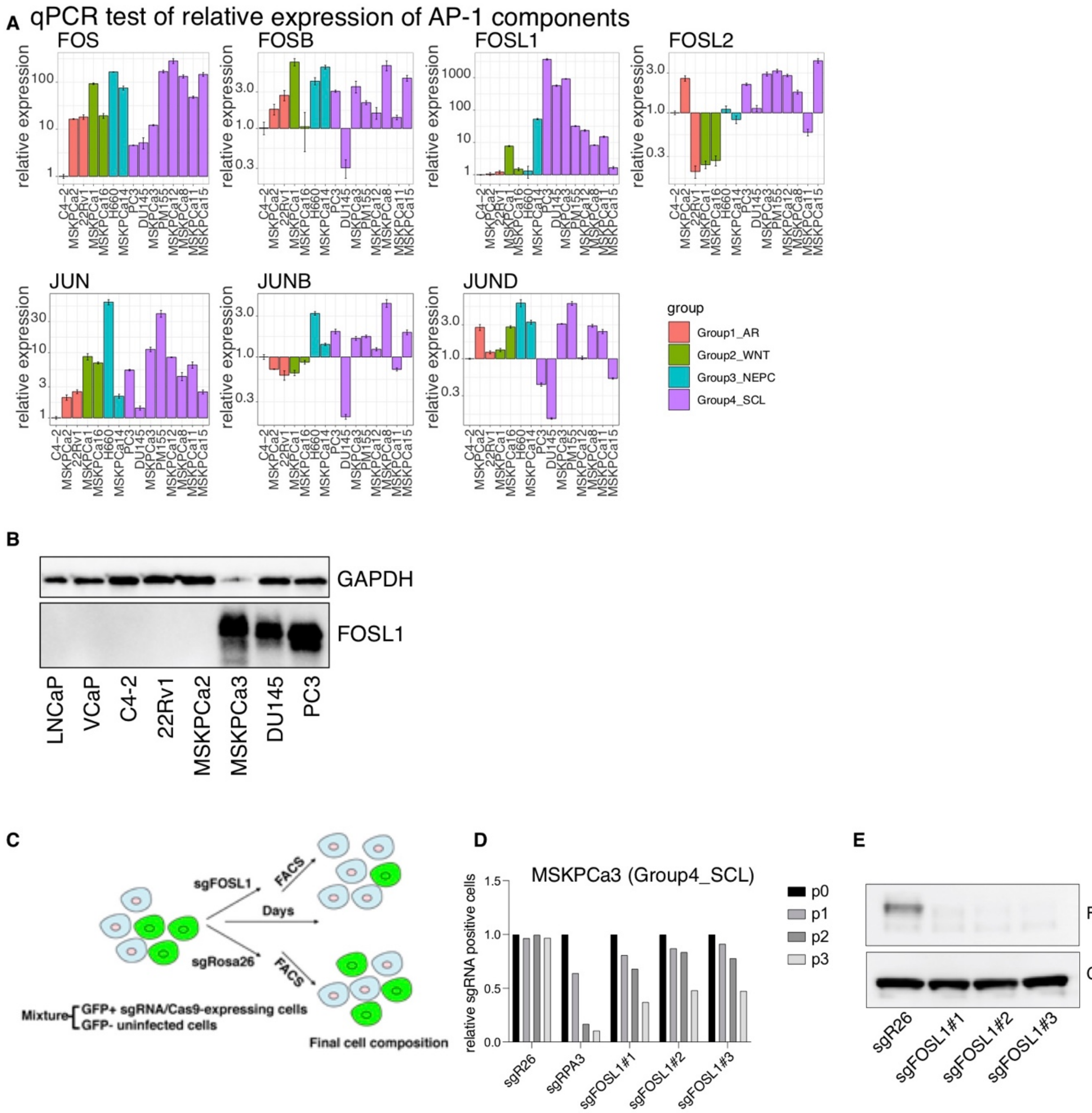

**Fig. S16 FOSL1 has higher relative expression and promotes tumor growth in Group4_SCL samples.** (**A**) qPCR analysis showing relative expression of FOS/JUN members across representative cell lines and organoids of the 4 subtypes. (**B**) Immunoblot indicating FOSL1 is not detectable in Group1_AR samples at the protein level. (**C**) Schematic showing experimental design for CRISPR competition assays. (**D**) Percentage of GFP positive MSKPCa3 expressing CRISPR guides against FOSL1 or sgRosa26 (negative control) or sgRPA3 (positive control) over 4 passages. (**E**) Knockout of FOSL1 was confirmed at protein level by immunoblot.

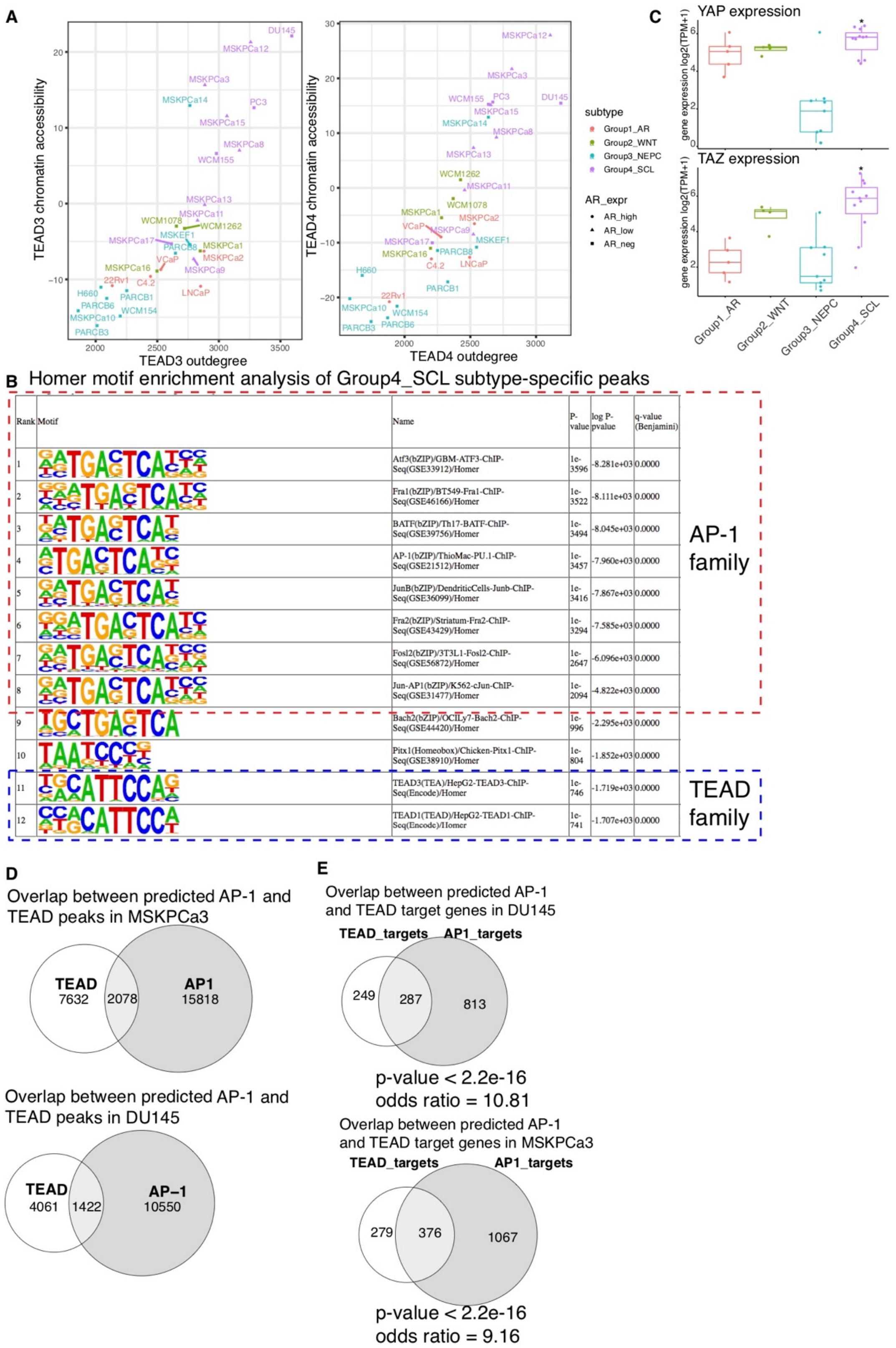

| Rank | Motif | Name | P-value | log P-pvalue | q-value (Benjamini) |
|---|---|---|---|---|---|
| 1 | | Atf3(bZIP)/GBM-ATF3-ChIP-Seq(GSE33912)/Homer | 1e-3596 | -8.281e+03 | 0.0000 |
| 2 | | Fra1(bZIP)/BT549-Fra1-ChIP-Seq(GSE46166)/Homer | 1e-3522 | -8.111e+03 | 0.0000 |
| 3 | | BATF(bZIP)/Th17-BATF-ChIP-Seq(GSE39756)/Homer | 1e-3494 | -8.045e+03 | 0.0000 |
| 4 | | AP-1(bZIP)/ThioMac-PU.1-ChIP-Seq(GSE21512)/Homer | 1e-3457 | -7.960e+03 | 0.0000 |
| 5 | | JunB(bZIP)/DendriticCells-Junb-ChIP-Seq(GSE36099)/Homer | 1e-3416 | -7.867e+03 | 0.0000 |
| 6 | | Fra2(bZIP)/Striatum-Fra2-ChIP-Seq(GSE43429)/Homer | 1e-3294 | -7.585e+03 | 0.0000 |
| 7 | | Fosl2(bZIP)/3T3L1-Fosl2-ChIP-Seq(GSE56872)/Homer | 1e-2647 | -6.096e+03 | 0.0000 |
| 8 | | Jun-AP1(bZIP)/K562-cJun-ChIP-Seq(GSE31477)/Homer | 1e-2094 | -4.822e+03 | 0.0000 |
| 9 | | Bach2(bZIP)/OCILy7-Bach2-ChIP-Seq(GSE44420)/Homer | 1e-996 | -2.295e+03 | 0.0000 |
| 10 | | Pitx1(Homeobox)/Chicken-Pitx1-ChIP-Seq(GSE38910)/Homer | 1e-804 | -1.852e+03 | 0.0000 |
| 11 | | TEAD3(TEA)/HepG2-TEAD3-ChIP-Seq(Encode)/Homer | 1e-746 | -1.719e+03 | 0.0000 |
| 12 | | TEAD1(TEAD)/HepG2-TEAD1-ChIP-Seq(Encode)/Homer | 1e-741 | -1.707e+03 | 0.0000 |

**Fig. S17 TEAD is enriched in Group4_SCL-specific ATAC-seq peaks, and functions together with AP-1.** (**A**) Group4_SCL are enriched with samples of high TEAD3/TEAD4 out-degree (from the constructed regulatory networks) and chromatin accessibility (z-score from ChromVAR). (**B**) Homer motif enrichment analysis indicates

Group4_SCL subtype-specific peaks are enriched with AP-1 and TEAD motifs. (**C**) Group4_SCL organoids and cell lines have higher expression of YAP and TAZ on mRNA level. The comparison is done between each group to the overall expression across all samples with one-sided Wilcoxon test with FDR correction. (**D**) Venn diagrams and permutation tests show significant overlaps are observed between predicted TEAD and AP-1 peaks in DU145 and MSKPCa3 (See table S11 for more details). (**E**) Fisher Exact Test shows the predicted TEAD and AP-1 targeted genes are significantly overlapped in DU145 and MSKPCa3.

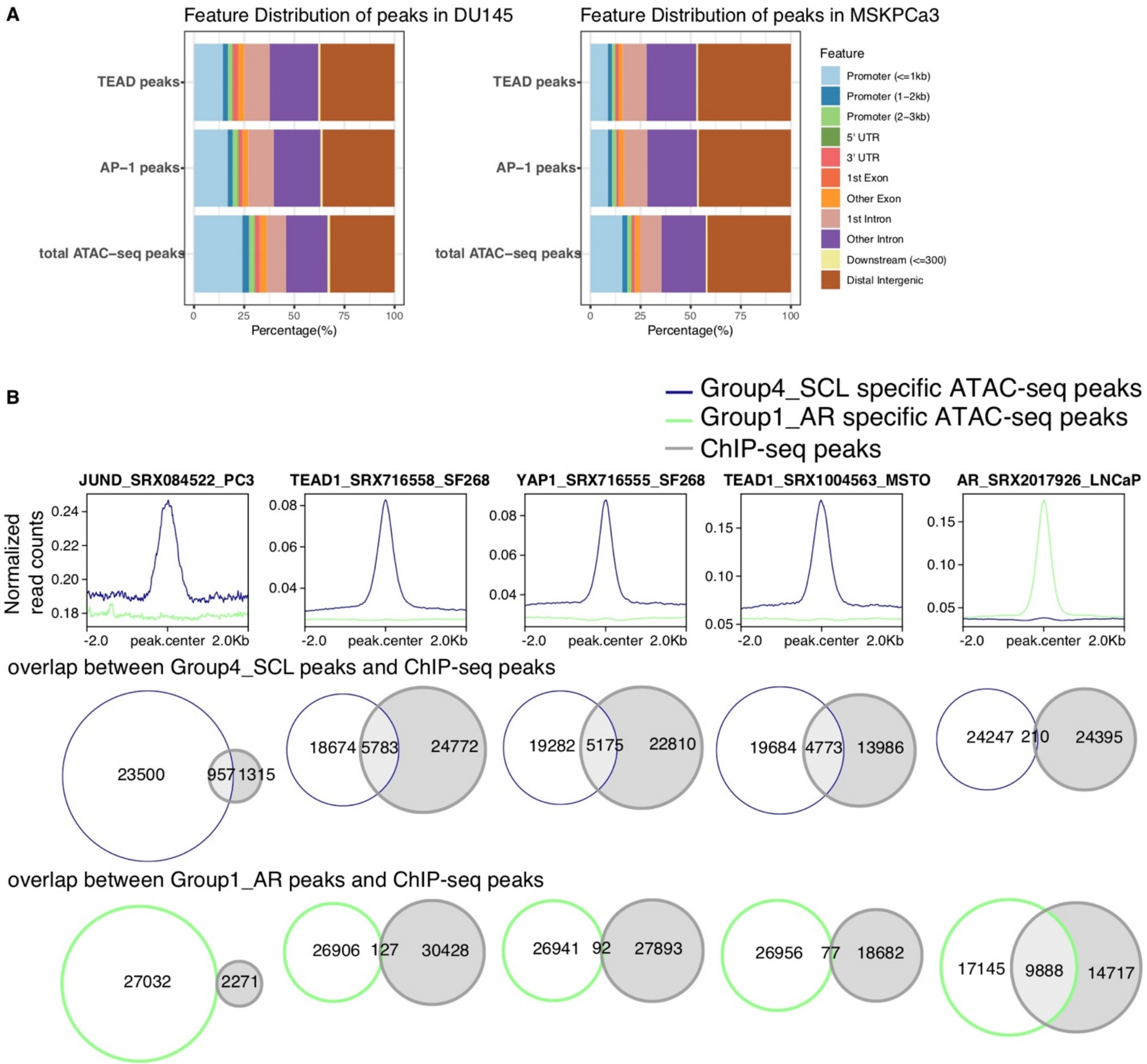

**Fig. S18 Additional ATAC-seq and ChIP-seq data suggests AP-1 works together with YAP, TAZ and TEAD.** (**A**) Feature distribution of the predicted TEAD, AP-1 and total ATAC-seq peaks in DU145 and MSKPCa3. (**B**) Similar to Fig5D. AP-1 (GSE29808), TEAD (GSE61852, GSE68170) and YAP (GSE61852) ChIP-seq peaks from multiple studies show stronger signal and more overlap with Group4_SCL specific ATAC-seq peaks while AR (GSE61852) ChIP-seq have stronger signal in Group1_AR specific peaks.

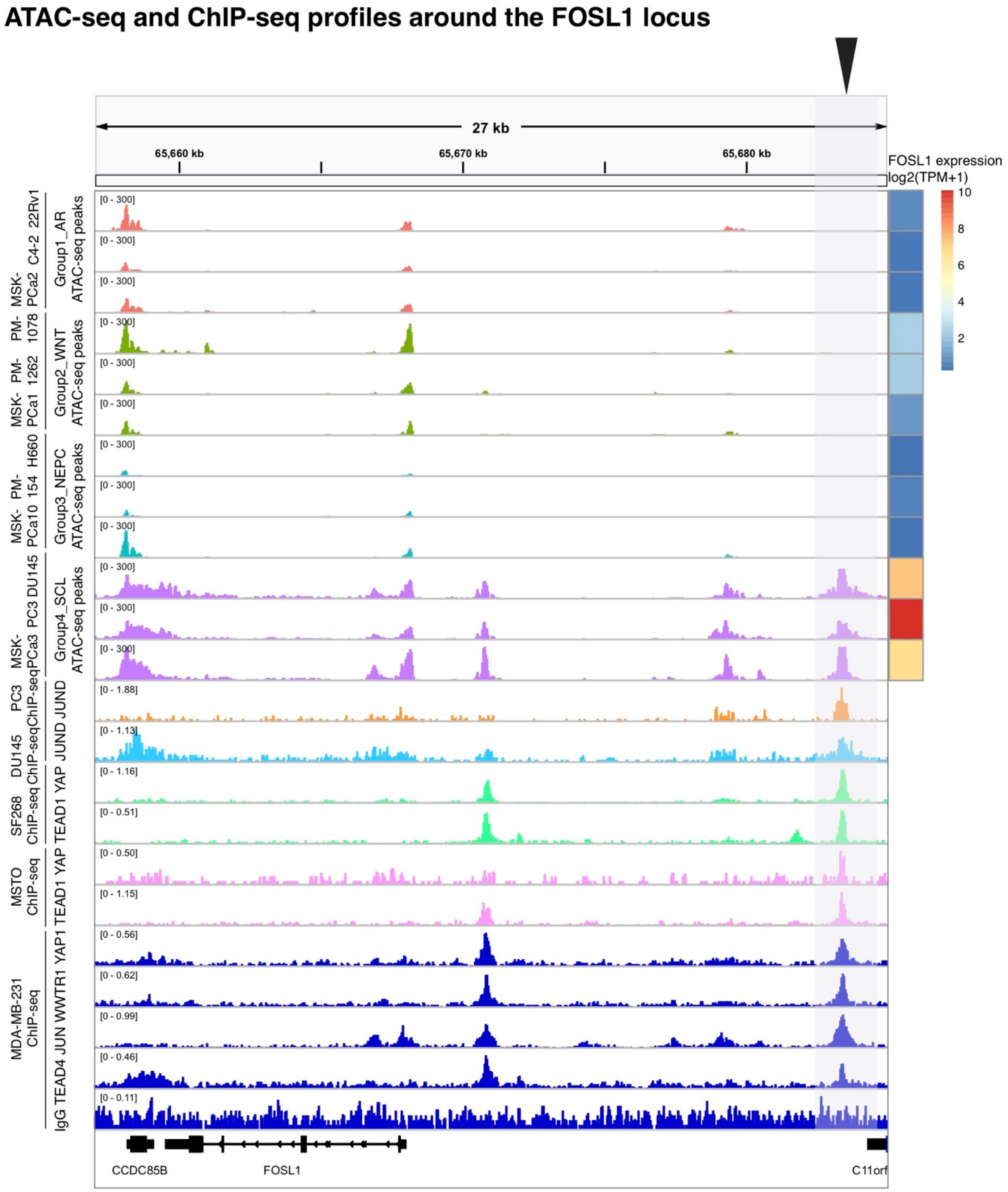

**Fig. S19 ATAC-seq and ChIP-seq profiles around the FOSL1 locus suggest that AP-1, YAP/TAZ and TEAD bind to the enhancer region of FOSL1 and increase its expression.** ATAC-seq peak tracks are kept to the same scale for comparison. On the right is the expression of FOSL1 across the representative organoids and cell lines from Group1_AR, Group2_WNT, Group3_NEPC and Group4_SCL. The highlighted ATAC-seq peaks are predicted to regulate FOSL1 expression (Table S6). The Gene Expression Omnibus (GEO) ID of the ChIP-seq peaks are listed in table S15.

### ATAC-seq and ChIP-seq profiles around the CYR61 locus

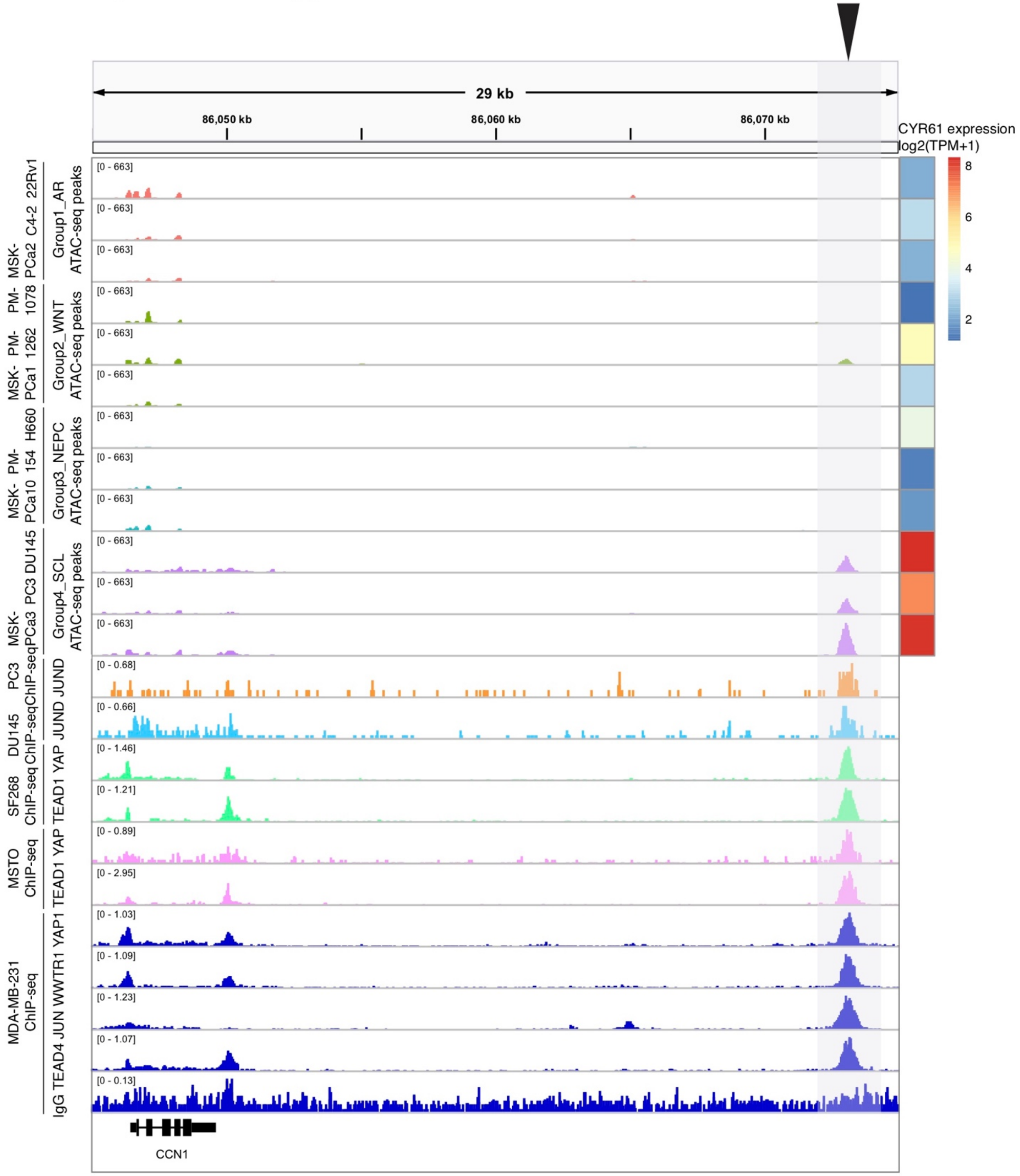

**Fig. S20 ATAC-seq and ChIP-seq profiles around the CYR61 locus suggest that AP-1, YAP/TAZ and TEAD bind to the enhancer region of CYR61 and increase its expression.** The highlighted ATAC-seq peaks are predicted to regulate CYR61 expression (Table S6). The GEO of the ChIP-seq peaks are the same as the ones used in Fig. S19.

### ATAC-seq and ChIP-seq profiles around the AXL locus

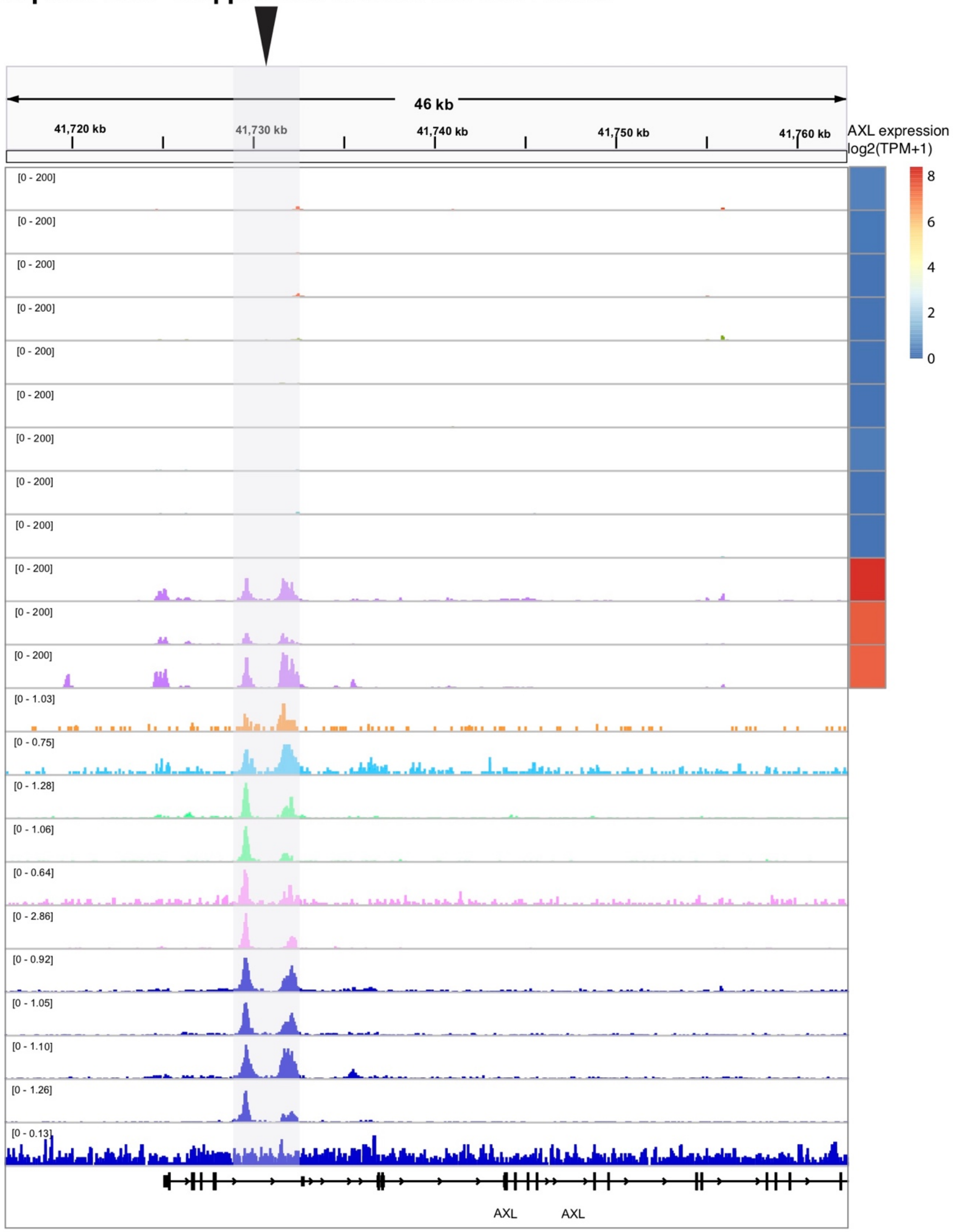

**Fig. S21 ATAC-seq and ChIP-seq profiles around the AXL locus suggest that AP-1, YAP/TAZ and TEAD bind to the gene body of AXL and increase its expression.** The highlighted ATAC-seq peaks are predicted to regulate AXL expression (Table S6). The GEO of the ChIP-seq peaks are the same as the ones used in Fig. S19.

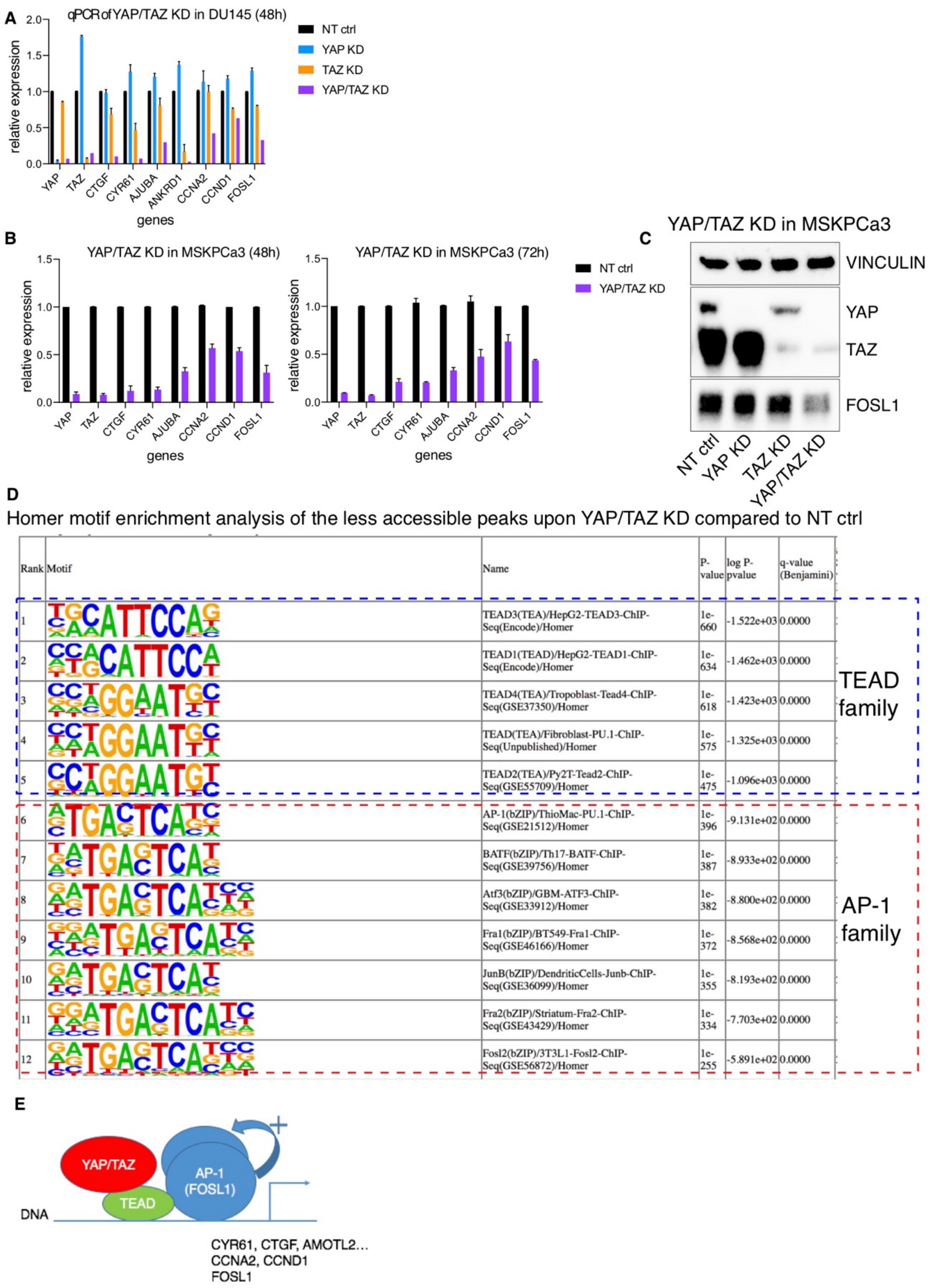

**Fig. S22 YAP/TAZ and AP-1 are essential for Group4_SCL samples DU145 and MSKPCa3.** (**A**) qPCR analysis result of select gene expression 48h after YAP/TAZ single or double knockdown of YAP/TAZ in DU145 (Group4_SCL). (**B**) qPCR of select gene expression 48h or 72h after YAP/TAZ double knockdown of YAP/TAZ in MSKPCa3 (Group4_SCL). (**C**) Immunoblot confirming the efficiency of YAP/TAZ knockdown by siRNA and the decrease of FOSL1 expression upon double knockout 72h after siRNA transfection in MSKPCa3. (**D**) Homer motif enrichment analysis indicates AP-1 and TEAD motifs are depleted upon YAP/TAZ KD. (**E**) Schematic illustrating the proposed model: YAP/TAZ work together with TEAD and AP-1 (FOSL1) to open chromatin and induce the expression of downstream genes, including canonical YAP/TAZ targets, cell cycle regulators and FOSL1 itself, forming a positive feedback loop.

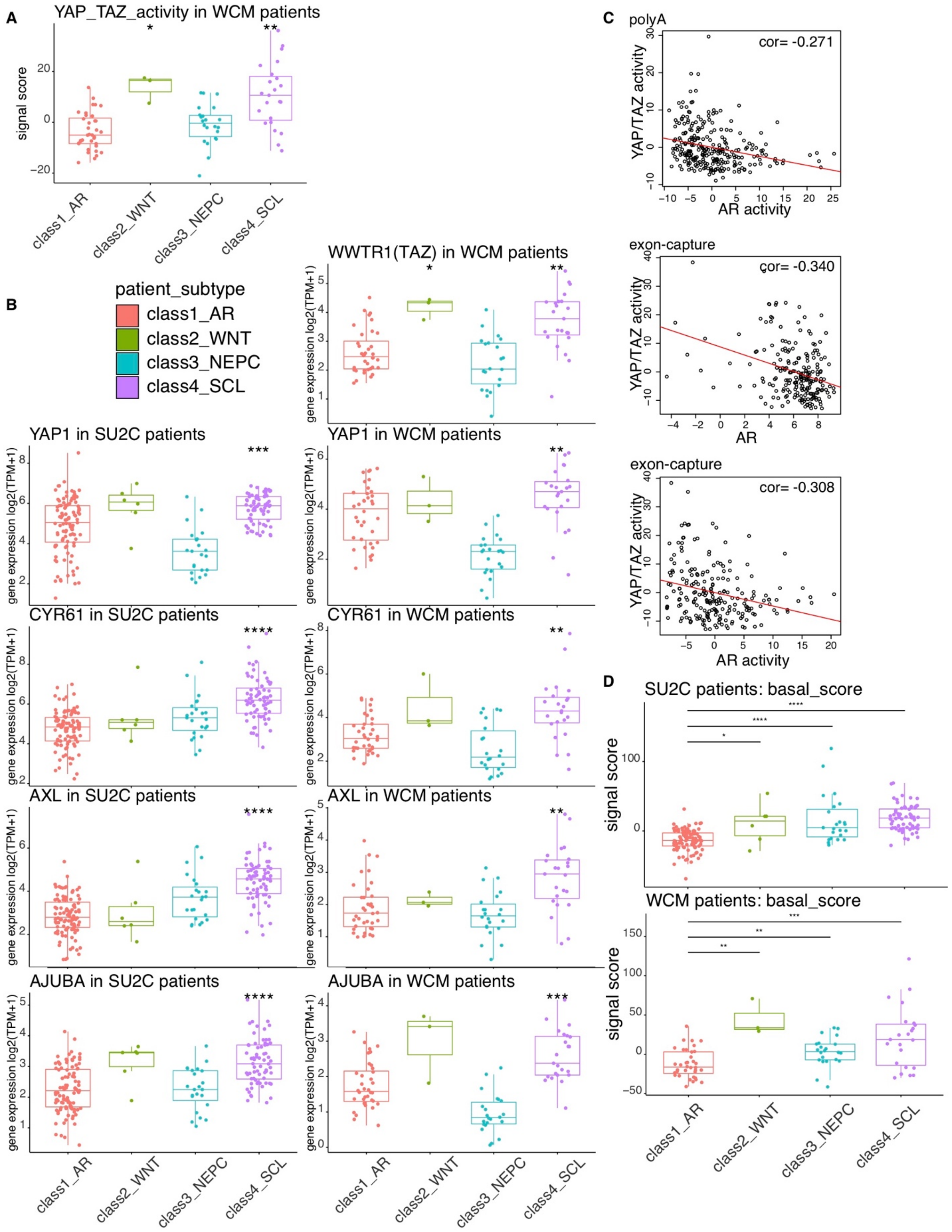

**Fig. S23 YAP/TAZ activity in patients.** (**A**) YAP/TAZ activity (sum of z-scores) is significantly higher in Group4_SCL WCM patients. (**B**) YAP, TAZ and selective downstream targets have higher expression in Group4_SCL patients. The comparison was performed between the scores of each group to the overall signal scores across all patients. (**C**) YAP/TAZ activity is significantly negatively correlated with AR expression and AR activity across 270 SU2C polyA assay samples and 212 exon-capture samples with $p < 0.001$. (**D**) Group2~Group4 patient tumor samples have higher basal score (sum of zscores) compared to Group1_AR patient samples for both SU2C (upper panel) and WCM (lower panel) cohorts. *$p<0.05$, **$p<0.01$, ***$p<0.001$, ****$p<0.0001$.

**Table S2.** Mutations detected by MSK-IMPACT (Memorial Sloan Kettering-Integrated Mutation Profiling of Actionable Cancer Targets) study.

| Biopsy | | | MSKPCa8 | | |
|---|---|---|---|---|---|
| **Tumor** | **AA change** | **AF** | **Organoids** | **AA change** | **AF** |
| *TP53* | L257P | 0.68 | *TP53* | L257P | 1 |
| *RB1* | F514Pfs*8 | 0.41 | *RB1* | F514Pfs*8 | 0.97 |
| *FOXA1* | H377Rfs*10 | 0.57 | *FOXA1* | H377Rfs*10 | 0.64 |
| *EGFR* | EGFR-intragenic fusion | n/a | | | |

| Biopsy | | | MSKPCa9 | | |
|---|---|---|---|---|---|
| **Tumor** | **AA change** | **AF** | **Organoids** | **AA change** | **AF** |
| *TP53* | I255F | 0.49 | *TP53* | I255F | 1 |
| *MED12* | R1357H | 0.69 | *MED12* | R1357H | 1 |
| | | | *SMAD4* | W524L | 1 |
| *CDH1* | Q673* | 0.29 | | | |
| *RB1* | RB1-intragenic fusion | n/a | | | |
| *EPHA5* | X836_splice | 0.13 | | | |

| Biopsy | | | MSKPCa10 | | |
|---|---|---|---|---|---|
| **Tumor** | **AA change** | **AF** | **Organoids** | **AA change** | **AF** |
| *TP53* | R282W | 0.55 | *TP53* | R282W | 0.53 |
| *TP53* | N239K | 0.32 | *TP53* | N239K | 0.49 |

| Biopsy | | | MSKPCa11 | | |
|---|---|---|---|---|---|
| **Tumor** | **AA change** | **AF** | **Organoids** | **AA change** | **AF** |
| *TP53* | Y103Tfs*20 | 0.58 | *TP53* | Y103Tfs*20 | 0.97 |
| *NOTCH1* | E1148* | 0.41 | *NOTCH1* | E1148* | 0.5 |
| *MDM4* | N390S | 0.57 | *MDM4* | N390S | 0.54 |

| Biopsy | | | MSKPCa12 | | |
|---|---|---|---|---|---|
| **Tumor** | **AA change** | **AF** | **Organoids** | **AA change** | **AF** |
| *TP53* | D393Gfs*78 | 0.13 | *TP53* | D393Gfs*78 | 0.92 |
| *PNRC1* | A165V | 0.09 | *PNRC1* | A165V | 0.68 |
| *PIK3R1* | PIK3R1-intragenic fusion | | *PIK3R1* | PIK3R1-intragenic fusion | |

| Biopsy | | | MSKPCa13 | | |
|---|---|---|---|---|---|
| **Tumor** | **AA change** | **AF** | **Organoids** | **AA change** | **AF** |
| *TP53* | R213* | 0.69 | *TP53* | R213* | 1 |
| *ERG* | TMPRSS2-ERG fusion | | *ERG* | TMPRSS2-ERG fusion | |
| *TMPRSS2* | TMPRSS2-ERG fusion | | *TMPRSS2* | TMPRSS2-ERG fusion | |

| Biopsy Tumor | AA change | AF | MSKPCa14 Organoids | AA change | AF |
|---|---|---|---|---|---|
| *ERG* | TMPRSS2-ERG fusion | | *ERG* | TMPRSS2-ERG fusion | |
| *TMPRSS2* | TMPRSS2-ERG fusion | | *TMPRSS2* | TMPRSS2-ERG fusion | |
| *ATM* | D1604V | 0.47 | *ATM* | D1604V | 0.52 |
| *NOTCH2* | R1260H | 0.48 | *NOTCH2* | R1260H | 0.49 |

* Truncation
Abbreviations: AA = amino acid; AF = allele frequency

**Table S3.** Unmatched mutational analysis for three organoids without normal control.

**MSKPCa15**

| Organoids | AA change | AF |
|---|---|---|
| *SPOP* | W131R | 0.73 |
| *TP53* | L257V | 1 |
| *MAP2K4* | V321E | 1 |
| *CDKN1B* | A121Sfs*3 | 1 |
| *FAT1* | N4267Ifs*84 | 0.19 |
| *JAK2* | N1108S | 0.62 |
| *JAK3* | E739Q | 0.24 |
| *ATR* | L227V | 0.35 |
| *CCND2* | R18G | 0.1 |
| *DIS3* | V793A | 0.98 |
| *DNAJB1* | V283F | 0.26 |
| *FAT1* | V3147G | 1 |
| *PTPRT* | L783Wfs*76 | 0.26 |
| *RAD54L* | R587W | 1 |
| *SPEN* | A2615V | 0.99 |
| *TSC2* | F1510del | 0.31 |

**MSKPCa16**

| Organoids | AA change | AF |
|---|---|---|
| *CTNNB1* | G34R | 0.27 |
| *TP53* | I195V | 0.6 |
| *TP53* | I195S | 0.6 |
| *TP53* | R213* | 0.33 |
| *TP53* | I195Mfs*13 | 0.61 |
| *APC* | R2326* | 0.55 |
| *PTPRS* | C27Y | 0.34 |
| *CREBBP* | A981T | 1 |
| *CREBBP* | V1129I | 1 |
| *FOXA1* | F254_L260del | 0.5 |
| *IRS2* | G1236R | 0.99 |
| *ERG* | R374C | 0.32 |
| *ERG* | R130C | 0.34 |
| *FGFR3* | A736Rfs*83 | 0.1 |
| *NCOR1* | V50M | 0.64 |
| *RPTOR* | D1151N | 0.36 |

**MSKPCa17**

| Organoids | AA change | AF |
|---|---|---|

| | | |
|---|---|---|
| *FLT3* | V194M | 0.98 |
| *RUNX1* | L56S | 0.49 |
| *INHA* | V163M | 0.3 |
| *PDGFRB* | P345S | 0.63 |
| *CASP8* | I357V | 0.69 |
| *CHEK2* | R449H | 0.66 |
| *ERBB2* | I654V | 0.47 |
| *NOTCH3* | H170R | 0.99 |
| *RAD50* | K446E | 0.39 |
| *ROS1* | T623S | 0.99 |
| *SH2B3* | S186I | 0.39 |

**Table S11.** The Fisher exact test results of overlapping between predicted TEAD-bound peaks and predicted master TF-bound peaks in representative samples in each of the four groups (related to fig. S17D)

| Sample | Group | TF (to overlap with TEAD) | p-value | odds ratio |
|---|---|---|---|---|
| DU145 | Group4_SCL | AP-1 | 1.85E-12 | 1.431705 |
| MSKPCa3 | Group4_SCL | AP-1 | 0.0001366 | 1.154953 |
| 22Rv1 | Group1_AR | AR | 0.4938 | 1.003534 |
| MSKPCa2 | Group1_AR | AR | 0.04009 | 1.100827 |
| PM1078 | Group2_WNT | TCF | 1 | 0.2222197 |
| MSKPCa16 | Group2_WNT | TCF | 1 | 0.1143611 |
| PM154 | Group3_NEPC | ASCL1 | 0.9961 | 0.8558703 |
| MSKPCa10 | Group3_NEPC | ASCL1 | 0.9668 | 0.8940321 |
| PM154 | Group3_NEPC | NEUROD1 | 0.9783 | 0.898862 |
| MSKPCa10 | Group3_NEPC | NEUROD1 | 1 | 0.7940421 |

**Table S12.** The antibody list used for immunoblot and IHC.

| **Antigen** | **Supplier** | **Ig type** | **Dilution** |
|---|---|---|---|
| AR | Abcam #ab108341 | Rabbit IgG | WB 1:2000 |
| FOSL1 | Cell Signaling #5281S | Rabbit IgG | WB 1:1000 |
| PTEN | Cell Signaling #9188 | Rabbit IgG | WB 1:1000 |
| RB1 | Cell Signaling #9309 | Mouse IgG | WB 1:1000 |
| SYP | Epitomics #1485-1 | Rabbit IgG | IHC 1:100 |
| SYP | Abcam #ab32127 | Rabbit IgG | WB 1:1000 |
| YAP/TAZ | Cell Signaling #8418S | Rabbit IgG | WB 1:1000 |
| GAPDH | abm #G041 | Mouse IgG | WB 1:10000 |
| VINCULIN | Abcam #ab129002 | Rabbit IgG | WB 1:10000 |

**Table S13.** The primer sequence for RT-qPCR.

| | |
|---|---|
| GAPDH_FW | CCAGGTGGTCTCCTCTGACTTC |
| GAPDH_RE | TCATACCAGGAAATGAGCTTGACA |
| RPL13A_FW | CCTGGAGGAGAAGAGGAAAGAGA |
| RPL13A_RE | TTGAGGACCTCTGTGTATTTGTCAA |
| TCF7L2_FP | GAATCGTCCCAGAGTGATGTC |
| TCF7L2_RP | ACGACCTTTGCTCTCATTTCC |
| AXIN2_FP | TGTCCAGCAAAACTCTGAGG |
| AXIN2_RP | GTGCAAAGACATAGCCAGAAC |
| RNF43_FP | GCTGGTGTTGCTGAAATAACTC |
| RNF43_RP | AATCCAGGCTCCAGATTGTC |
| FOSL1_FP | AGGCCTTGTGAACAGATCAG |
| FOSL1_RP | TCATCTTCCAGTTTGTCAGTCTC |
| FOSL2_FP | CGGGAGCTGACAGAGAAG |
| FOSL2_RP | GGGCTAATCTTGCACACTGG |
| FOS_FW | TTGTGAAGACCATGACAGGAG |
| FOS_RE | CCATCTTATTCCTTTCCCTTCGG |
| FOSB_FW | AGCTAAATGCAGGAACCGG |
| FOSB_RE | ACCAGCACAAACTCCAGAC |
| JUND_FW | ATCGACATGGACACGCAG |
| JUND_RE | GGGTCTTCACTTTCTCTTCCAG |
| JUNB_FP | GGACACGCCTTCTGAACG |
| JUNB_RP | CGGAGTCCAGTGTGGTTTG |
| JUN_FW | AGCCCAAACTAACCTCACG |
| JUN_RE | TGCTCTGTTTCAGGATCTTGG |
| ASCL1_FW | CAACGCCACTGACAAGAAAG |
| ASCL1_RE | GGAGCTTCTCGACTTCACCA |
| NKX2-1_FW | AACCAAGCGCATCCAATCTCAAGG |
| NKX2-1_RE | TGTGCCCAGAGTGAAGTTTGGTCT |
| NEUROD1_FW | CCAGGGTTATGAGACTATCACTG |
| NEUROD1_RE | TCCTGAGAACTGAGACACTCG |
| TMPRSS2_FW | GGACAGTGTGCACCTCAAAGAC |
| TMPRSS2_RE | TCCCACGAGGAAGGTCCC |
| CCNA2_FW | GTGAAGATGCCCTGGCTTTTA |
| CCNA2_RE | TCCTCATGGTAGTCTGGTACTT |
| CCND1_FW | GCTGCGAAGTGGAAACCATCC |
| CCND1_RE | CGCACTTCTGTTCCTCGCAG |
| WWTR1_FW | AGCTCAGATCCTTTCCTCAATG |
| WWTR1_RE | TCCTGCGTTTTCTCCTGTATC |

| | |
|---|---|
| YAP1_FW | AGATGGAGAAGGAGAGGCTG |
| YAP1_RE | AGTGTTGGTAACTGGCTACG |
| AJUBA_FW | TGTCAATGGCTCTGTGTACTG |
| AJUBA_RE | ACTTCCCCATTGCTTGTAGG |
| ANKRD1_FW | GGTGAGACTGAACCGCTATAAG |
| ANKRD1_RE | GGCTGTCGAATATTGCTTTGG |
| CYR61_FW | CAAGGAGCTGGGATTCGATG |
| CYR61_RE | AAAGGGTTGTATAGGATGCGAG |
| CTGF_FW | TGCCCTCGCGGCTTACCGACTG |
| CTGF_RE | TGCAGGAGGCGTTGTCATTGGTAAC |
| AMOTL2_FW | AAAGCAGGTTAAAGGTGCTCCA |
| AMOTL2_RE | CTCTGTGGGTGCTCTGTCTG |

**Table S15.** The GEO ID of the public available data used in the paper.

| **RNA-seq** | | | |
|---|---|---|---|
| **sample** | **GSE** | **ref paper** | |
| MSKEF1 | GSE118207 | https://pubmed.ncbi.nlm.nih.gov/30287662/ | |
| PARCB1 | GSE118207 | https://pubmed.ncbi.nlm.nih.gov/30287662/ | |
| PARCB3 | GSE118207 | https://pubmed.ncbi.nlm.nih.gov/30287662/ | |
| PARCB6 | GSE118207 | https://pubmed.ncbi.nlm.nih.gov/30287662/ | |
| PARCB8 | GSE118207 | https://pubmed.ncbi.nlm.nih.gov/30287662/ | |
| 22Rv1 | GSE118207 | https://pubmed.ncbi.nlm.nih.gov/30287662/ | |
| LNCaP | GSE118207 | https://pubmed.ncbi.nlm.nih.gov/30287662/ | |
| VCaP | GSE118207 | https://pubmed.ncbi.nlm.nih.gov/30287662/ | |
| DU145 | GSE118207 | https://pubmed.ncbi.nlm.nih.gov/30287662/ | |
| H660 | GSE118207 | https://pubmed.ncbi.nlm.nih.gov/30287662/ | |
| C4-2 | GSE114600 | https://pubmed.ncbi.nlm.nih.gov/30380416/ | |
| **ATAC-seq** | | | |
| **sample** | **GSE** | **ref paper** | |
| MSKEF1 | GSE118207 | https://pubmed.ncbi.nlm.nih.gov/30287662/ | |
| PARCB1 | GSE118207 | https://pubmed.ncbi.nlm.nih.gov/30287662/ | |
| PARCB3 | GSE118207 | https://pubmed.ncbi.nlm.nih.gov/30287662/ | |
| PARCB6 | GSE118207 | https://pubmed.ncbi.nlm.nih.gov/30287662/ | |
| PARCB8 | GSE118207 | https://pubmed.ncbi.nlm.nih.gov/30287662/ | |
| **ChIP-seq** | | | |
| **sample** | **GSE** | **ref paper** | **ChIP TF** |
| LNCaP | GSE52725 | https://pubmed.ncbi.nlm.nih.gov/24423874/ | AR |
| LNCaP | GSE85559 | https://pubmed.ncbi.nlm.nih.gov/29153843/ | AR |
| MDA-MB-231 | GSE66083 | https://pubmed.ncbi.nlm.nih.gov/26258633/ | JUN |
| MDA-MB-231 | GSE66083 | https://pubmed.ncbi.nlm.nih.gov/26258633/ | TEAD4 |
| MDA-MB-231 | GSE66083 | https://pubmed.ncbi.nlm.nih.gov/26258633/ | YAP1 |
| MDA-MB-231 | GSE66083 | https://pubmed.ncbi.nlm.nih.gov/26258633/ | WWTR1 |
| MDA-MB-231 | GSE66083 | https://pubmed.ncbi.nlm.nih.gov/26258633/ | IgG |
| SF268 | GSE61852 | https://pubmed.ncbi.nlm.nih.gov/26295846 | TEAD1 |
| SF268 | GSE61852 | https://pubmed.ncbi.nlm.nih.gov/26295846 | YAP1 |
| MSTO | GSE68170 | https://pubmed.ncbi.nlm.nih.gov/26439301/ | TEAD1 |
| MSTO | GSE68170 | https://pubmed.ncbi.nlm.nih.gov/26439301/ | YAP1 |
| PC3 | GSE29808 | https://pubmed.ncbi.nlm.nih.gov/22012618/ | JUND |
| DU145 | GSE59021 | https://pubmed.ncbi.nlm.nih.gov/25294825/ | JUND |

**Captions for Tables S1, S4 to S11, S15:**

**Table S1. (separate file)**
Patient and Organoid Characteristics.

**Table S4. (separate file)**
Sequencing statistics and ATAC-seq QC

**Table S5. (separate file)**
The RB1, TP53 and PTEN status of the 29 organoids and cell lines.

**Table S6. (separate file)**
Predicted peak-to-gene links.

**Table S7. (separate file)**
The TF_ranks for each of the four subtypes.

**Table S8. (separate file)**
The signature genes for SU2C and WCM patient classification.

**Table S9. (separate file)**
The SU2C and WCM patient classification result from nearest template prediction.

**Table S10. (separate file)**
Overall survival time and time on first-line ARSI for SU2C patients classified into the four subtypes.

**Table S14. (separate file)**
The gene set enrichment analysis signatures used in the paper.
